# Multiplexed analysis of a single EV (MASEV) reveals unique biomarker composition with diagnostic impact

**DOI:** 10.1101/2022.06.17.496607

**Authors:** Joshua Spitzberg, Scott Ferguson, Katy Yang, Hannah M. Peterson, Jonathan C.T. Carlson, Ralph Weissleder

## Abstract

Exosomes and extracellular vesicles (EV) are increasingly being explored as circulating biomarkers, but their heterogenous composition will likely mandate the development of single EV technologies. Highly multiplexed analyses of single EVs have been challenging to implement beyond a few colors during spectral sensing. We use a multiplexed analysis of the single EV technique (MASEV) to interrogate thousands of individual EVs during 5 cycles of multi-channel fluorescence staining for 15 EV biomarkers. Contrary to the common belief, we show that i) several “ubiquitous” markers are much less common than believed; ii) that multiple biomarkers concur in single vesicles but only in small fractions, iii) that affinity purification can lead to loss of rare EV subtypes, and iv) that deep profiling allows detailed analysis of EV potentially improving the diagnostic content. These findings establish the potential of MASEV for uncovering fundamental EV biology and heterogeneity and increasing diagnostic specificity.

**Graphical abstract.**
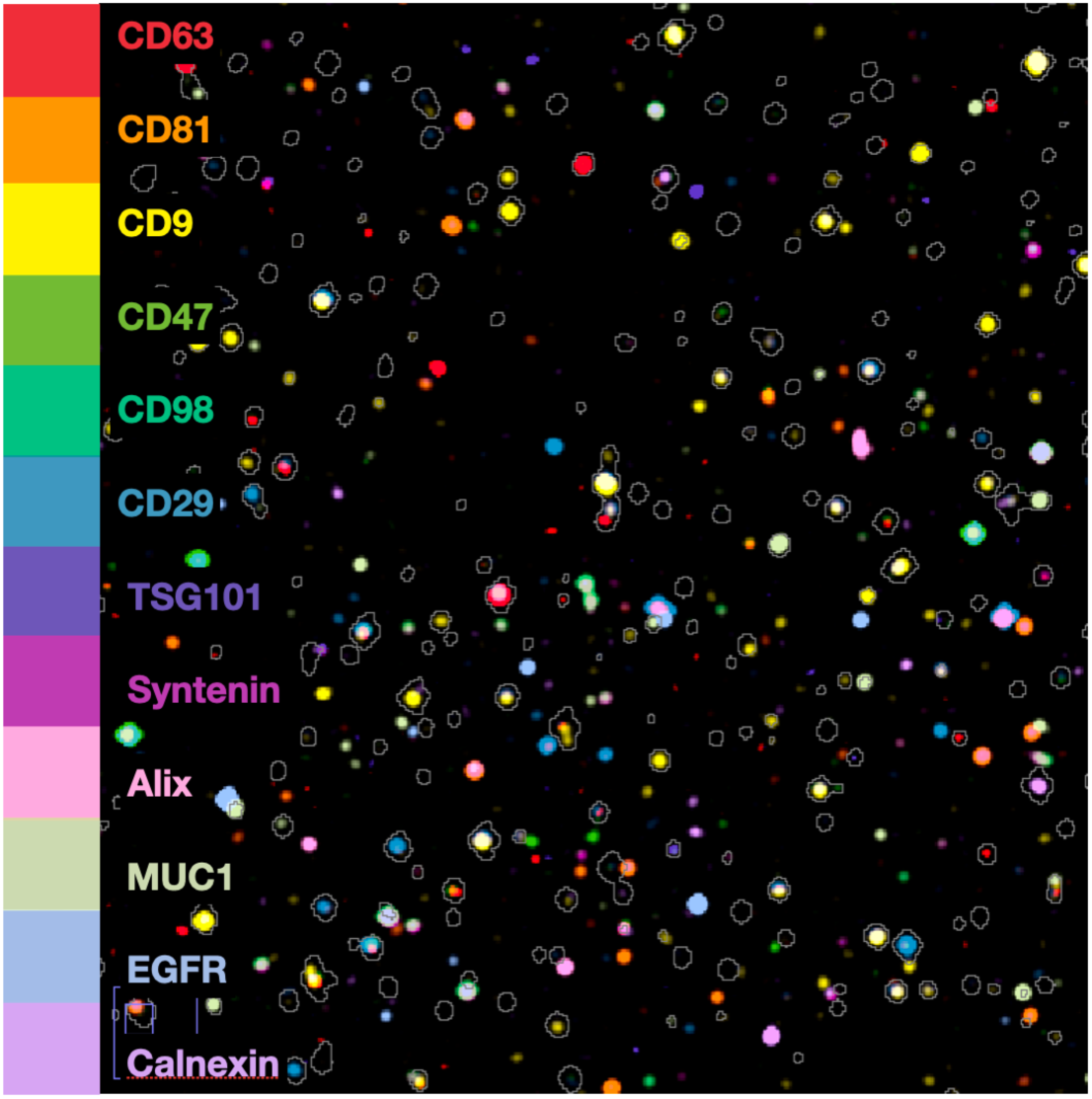
Multiplexed analysis of single EV (MASEV) allows robust protein profiling at the single EV level, a prerequisite for early cancer detection or organ of origin determination.

## Introduction

There is substantial interest in liquid biopsy approaches for cancer care, including early detection^1^. In particular, circulating tumor-derived extracellular vesicles (EV) represent a promising venue because shed vesicles are stable, contain cargo derived from parental cells^2, 3^, and are abundant in later-stage disease. Similar to other biomarker types (e.g., ctDNA^4, 5^, circulating tumor cells (CTC)^6^, proteins^7, 8^, metabolites^9^), the challenge in early cancers is to i) improve detection sensitivities of existing technologies, ii) define tumor-specific mutations and biomarkers, iii) differentiate tumor cell from host cell-derived vesicles with confidence, and iv) develop clinically viable technologies that can be tested in prospective trials. Technological advances have improved our ability to isolate and analyze bulk EV in plasma and biofluids. Recent advances are in part due to the miniaturization of detection technology^10^, integrated sensor platforms^11^ capable of point-of-care testing in a clinical environment^12^, digital sensing approaches^13^, amplification strategies^12^ and consensus on pre-analytical purification^10, 14–17^.

An important and essential advance to EV profiling has been the introduction of single EV (sEV) analytical techniques such as SEA^18^. Various permutations have been reported over the last few years. For example, sEVA^19^ is an advancement over SEA as it does not require EV capture prior to staining but initiates profiling in the solution phase. Several other approaches^20–24^ are helpful research tools but perhaps too complex for routine clinical use. Irrespective of the specific sEV analytical technique, obtaining multiplexed data from single vesicles has remained challenging. However, this will likely be essential in defining vesicle subpopulations and identifying rare cancer-specific phenotypes early in the disease.

We hypothesized that recent advances in bioorthogonal chemistry^25, 26^ could be used to develop more efficient sEV multiplexing tools. Here we report on an innovative tetrazine/ *trans*-cyclooctene (Tz/TCO) scission approach^26^ to perform cycling on repetitively labeled single EV in a simple flow chamber (**Fig. S1**). Combined with the multichannel acquisition, we show that this technique (MASEV, multiplexed analysis of single EV) allows rapid profiling of ∼ 15 different markers in a single EV. We use MASEV to shed light on EV biomarker abundance and develop more resilient EV approaches for early cancer detection. Furthermore, we show that “ubiquitous” exosome biomarkers are often present in fewer than 30% of all EV in cell line samples.

## Results

### Bioorthogonal scission chemistry allows multiplexed analysis of a single EV (MASEV)

The MASEV technology employs a specialty linker between an antibody of interest and a fluorochrome. This linker contains a C_2_-symmetric TCO moiety (C_2_TCO)^26^, which is stable and does not affect the fluorescent properties of the affinity ligand (**Fig. S2**).

However, upon the addition of functionalized tetrazine scissors (HK-Tz), the fluorochrome is selectively cleaved from the antibody, leading to very fast and clean “de-staining” that removes >99% of the fluorophore-derived signal at exceptionally gentle micromolar concentrations (**Fig. 1**). Over 90% de-staining is complete within 1-2 minutes at these reaction kinetics, and EV can then be stained for subsequent rounds. The cyclic staining chemistry is also compatible with live cells and tissues^27^ and does not affect the properties of EV. We developed a specially constructed flow chamber to facilitate rapid cycling and preservation of reagents, with no need for harsh conditions (e.g., paraformaldehyde to fix and adhere EVs or hydrogen peroxide to bleach fluorochromes), thus allowing unequivocal analyses of a single EV. This chamber uses an acrylic pressure-sensitive adhesive to bond a treated coverglass to a microscope slide^28^. The shape of the pressure-sensitive adhesive (width 4 mm, length 12 mm, height 50 µm) allows pump-free flushing with flow rates of ∼ 1 µL/s within the ∼2 µL channel. The hydrophobic silanization treatment adheres EV and prevents the flowcell reservoir from leaking or wetting out over the duration of multiple staining rounds.

**Fig. 1:**
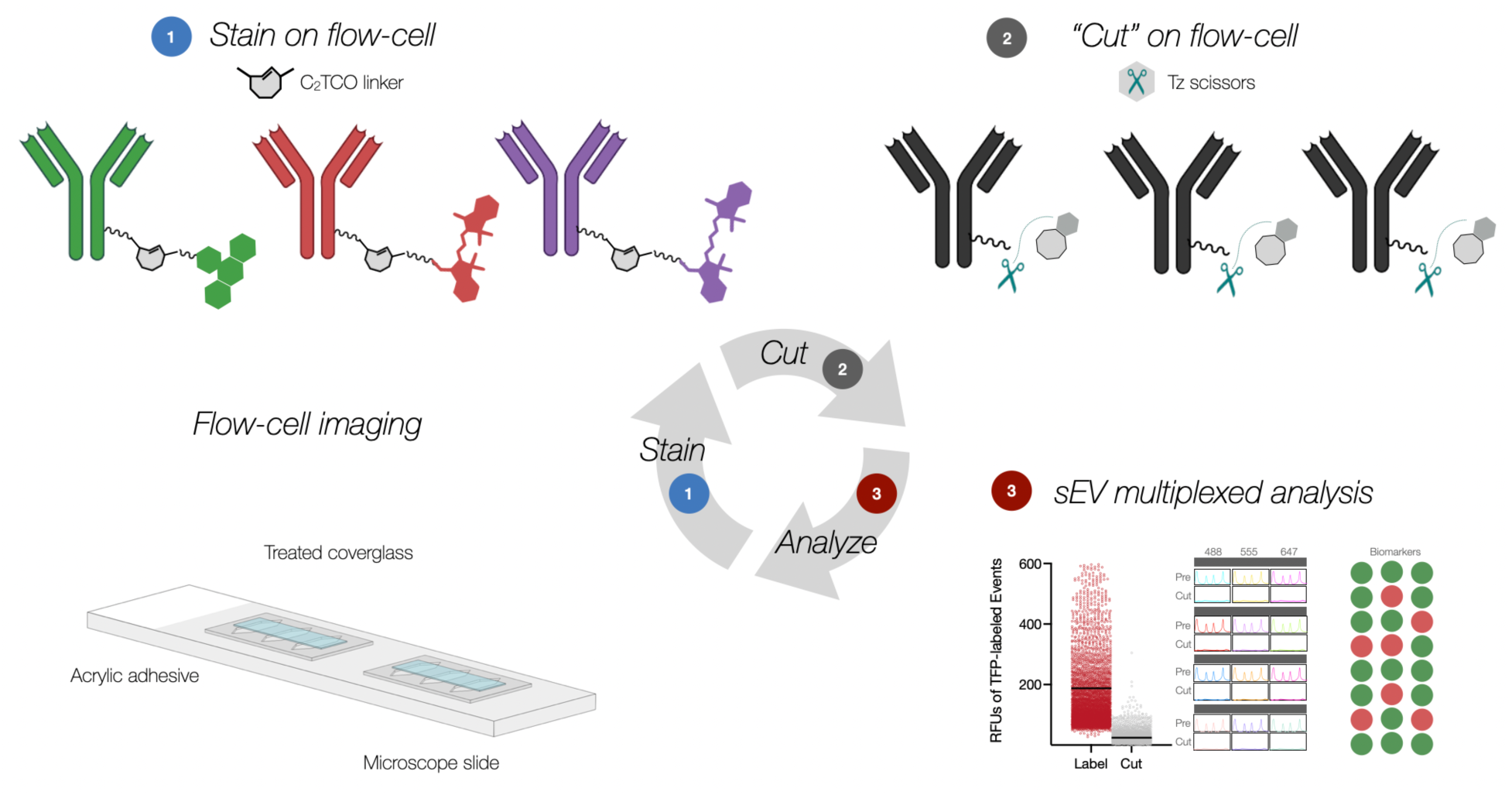
Overview of multiplexed analysis of single EV (MASEV). 1. Using a flow cell, pre-purified EV (labeled with unquenchable TFP-AF350) are attached to clean glass surfaces. Integrated channels allow uniform fluid flow at low pressure for subsequent staining, quenching, and washing cycles. EVs are first stained with up to three different fluorescently labeled antibodies, whereas the fourth TFP channel serves as a pan-EV reference. Fluorochromes are attached to antibodies via specialty linkers containing a C_2_TCO. **2**. Following image acquisition, the fluorochromes are cut by adding a tetrazine (Tz). This results in rapid and complete de-staining of all EVs in seconds. **3**. Fluorescence intensities before and after Tz addition are quantitated and plotted for each cycle. This data can generate biomarker profiles for a single EV using calibration curves.

Incubating the treated glass devices with Tween-20 reduced nonspecific antibody adhesion^29^, thus reducing background several fold relative to substrates such as plain glass slides (1.8x), ready-made adhesive-coated PTFE well-slides (1.8x), or poly-L-lysine coated slides (5.5x), each prepared with routine blocking buffers (e.g. SuperBlock).

### Optimization of MASEV

In the first set of experiments, we determined the retention rate of AlexaFluor350-PEG_12_-tetrafluorophenol (TFP_350_) labeled EV on glass slides^19^. This bright hydrophilic covalent marker reacts with free bioamines to provide a universal reference stain. Unlike prior studies^18^, we decided against covalent surface attachment because this approach can lead to selective loss of unbound EV and high background due to the chemistries involved. Therefore, we used extra clean cover glasses activated with KOH, functionalized with dichlorodimethylsilane, and (after incubating with EVs) blocked with Tween-20. These cover glasses, acrylic tape, and glass slides were used to construct mini chambers where Laplacian pressure-driven fluid flow was used to gently stain glass-adherent EVs (**Fig. S1** and **Movie 1**). Microscope slides were cleaned but not silanized because a hydrophobic base slide would reject liquid from its reservoirs and rapidly dry out the flow cell.

The data show that same-day experiments resulted in more than 99% EV retention across multiple cycles (**Fig 2A**). If the processes were protracted over two days, there was a 5 to 10% loss of EV on the second day. We, therefore, chose to perform all experiments in a single day. We also conducted several pre-analytical experiments to determine which EV preparations were most helpful. Specifically, we compared ultracentrifugation, traditional^17^, and dual-mode size exclusion chromatography (eDMC)^30^ to identify practical methods that yielded high amounts of EV. Our results show that the enhanced dual size-exclusion mode yielded high quantities of EV, was efficient, cost-effective, and allowed processing small sample sizes (**Fig. S3**). In summary, these results confirm work by others^14, 31–39^ that the size exclusion column methods of isolation yield pure EV for MASEV analysis in reasonable amounts of time.

**Fig. 2:**
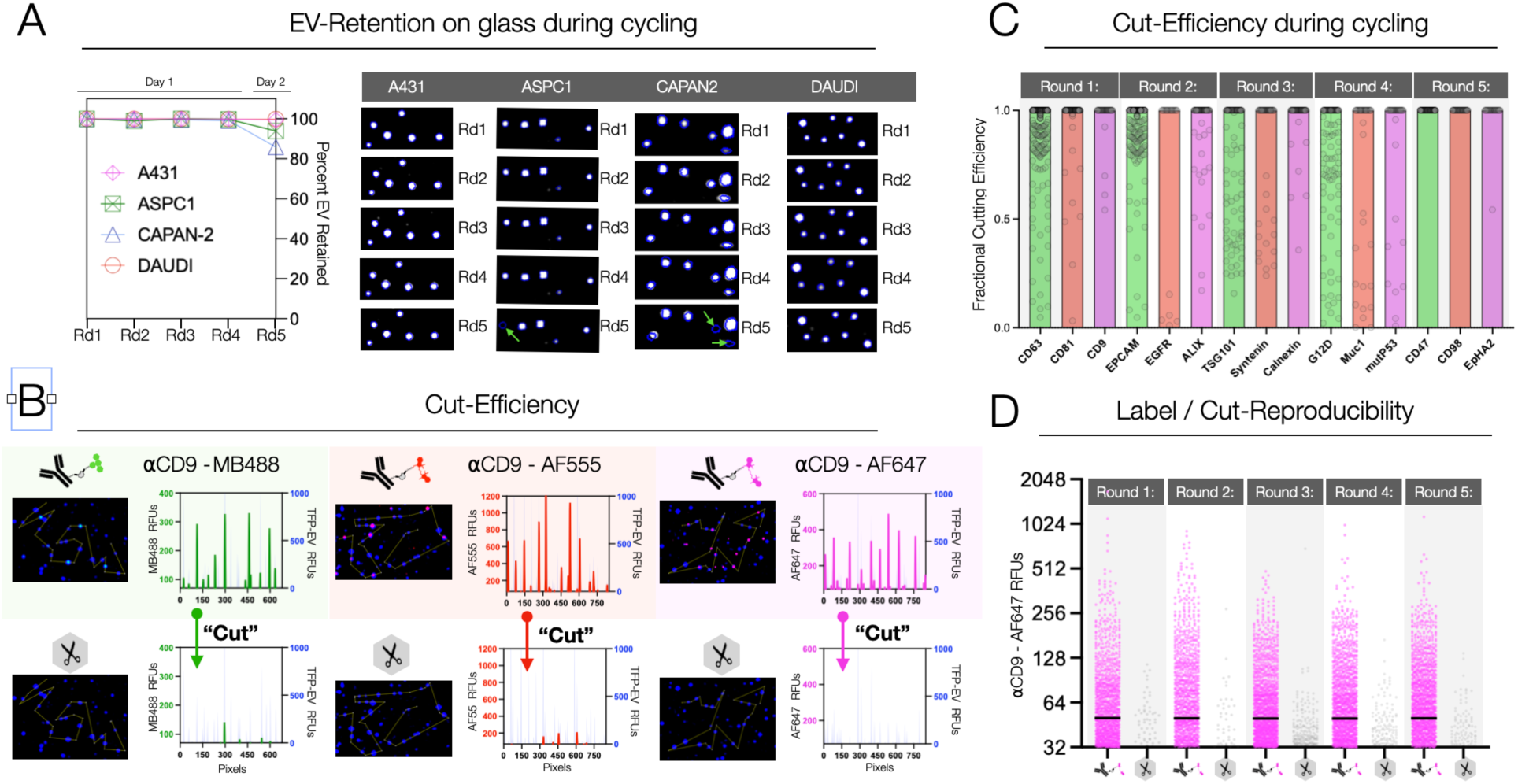
Characterization of the MASEV method. **A**. Retention of EV glass slide chambers. Note that the vast majority of EVs (> 99%) remain at their exact origin during four cycles of staining on day one but decrease slightly if experiments are spread out over two days (90% retention). **B.** Graphs demonstrate EV signal intensity after CD9 immunostaining and quenching for three fluorochromes: MB488, AF555, and AF647. Note the high SNR when labeled. **C.** Cut-efficiency of all antibodies. The cut-efficiency is > 99% (i.e., complete de-quenching of all EV), but the standard deviations are slightly higher with some antibodies. Since pre and post-cut data is available for each cycle, only EV with complete quenching (> 98%) are used for subsequent quantitative analysis (although all raw data is shown here). **D.** Reproducibility of immunostaining during different cycles. PANC1 EVs were repeatedly stained for CD9 across five cycles in this experiment. Note the nearly identical staining and de-staining during each process.

We next examined the on-chip signal dynamics and cycling efficiency of fluorophore addition and removal for single EV. Using CD9 as a prototypical EV marker, we processed EV from PANC-1 cell lines and measured signal intensities of stained versus de-stained EV with three different antibody-C_2_TCO-conjugated fluorochromes (MB488, AF555, or AF647) (**Fig. 2B**). The single-EV signal-to-background ratio (S/B) in the 488 channel was 5.7±0.4 for the stained EV, which dropped to 1.1±1.2 after destaining, with the mean EV signal indistinguishable from the slide background. Likewise, for the 555 and 647 channels, stained (vs. de-stained) S/B were 7.4±1.8 (vs. 1.2±0.4) and 6.0±0.4 (vs. 1.1±1.1), respectively. These values are consistent with the >95% scission observed in other contexts^26, 27^.

Next, we performed staining and de-staining experiments for an expanded panel of EV biomarkers (**Fig. 2C**). We used triplets of antibodies in a total of 5 rounds of staining and de-staining. The data shows that the cutting efficiency was 99% for the majority of C_2_TCO-antibody-labeled EVs across all three fluorescence channels. The lowest background was seen in the magenta channel (628/692 nm ex/em), while the red (562/593 nm) and green (472/520 nm) channels showed higher backgrounds (varying between 1.4-4 x compared to DAPI channel). Therefore, the judicious assignment of abundant - hence brighter - markers to the green and red channels allowed the processing of 15 biomarkers across EVs with minimal interference from background signal. In another set of experiments, we determined the reproducibility of staining and de-staining across the five cycles. As shown in **Fig. 2D**, the staining pattern was remarkably reproducible across the five cycles using CD9-AF647 as a prototypical marker. Indeed, the percent positive EV ranged from 22.5% to 27.6% across all rounds, with no evidence of background accumulation across the five cycles of destaining.

Finally, we performed additional validation experiments comparing the MASEV method to immunogold labeling for electron microscopy. Collectively, these data confirm that i) all labeled structures were indeed vesicles and ii) that the MASEV staining patterns correlated with immunogold labeling for two prototypical makers: CD63 and KRAS^mut^ (**Fig. S7**).

### Ubiquitous EV biomarkers are less common than thought

Bulk methods (Western, ELISA) often report tetraspanins, ALIX, TSG integrins, and syntenin as abundant and defining biomarkers necessary for EV formation^40^. Some other exosomal proteins have recently been found to be abundant, primarily through mass spectrometry analyses of bulk EV (CD47, CD29(ITGB1), ATP1A1, SLC1A5, SLC3A2, BSG)^33^. At the current time, what is unclear is how some of these putatively ubiquitous biomarkers are expressed on individual EVs. Are they all present at varying concentrations in all EV, or are some EV enriched in specific proteins? Defining such patterns could be helpful in future multiplexed analyses.

Starting with pure EV preparations obtained from PANC1 cells, we measured a 5-cycle, 15-plex panel (**Fig. 3**) to determine i) the abundance of each protein in a single EV and ii) the concurrence of multiple markers in individual vesicles. **Fig 3A** shows one representative example of such an experiment. As is evident from visual inspection of the images, there is a heterogeneous distribution of individual biomarkers across any EV population. In PANC1, the most abundant biomarkers were CD9 (47.9% of all EV), CD29 (26.0%), CD47 (19.9%), CD63 (12.9%), CD98 (11.5%), CD81 (7.3%) and Alix (6.5%). All other markers were present in fewer than 20% of EVs. Similar findings were observed across EV derived from other cell lines, with some variability in biomarker expression (**Fig. 3B, Fig S5-S6**).

**Fig. 3:**
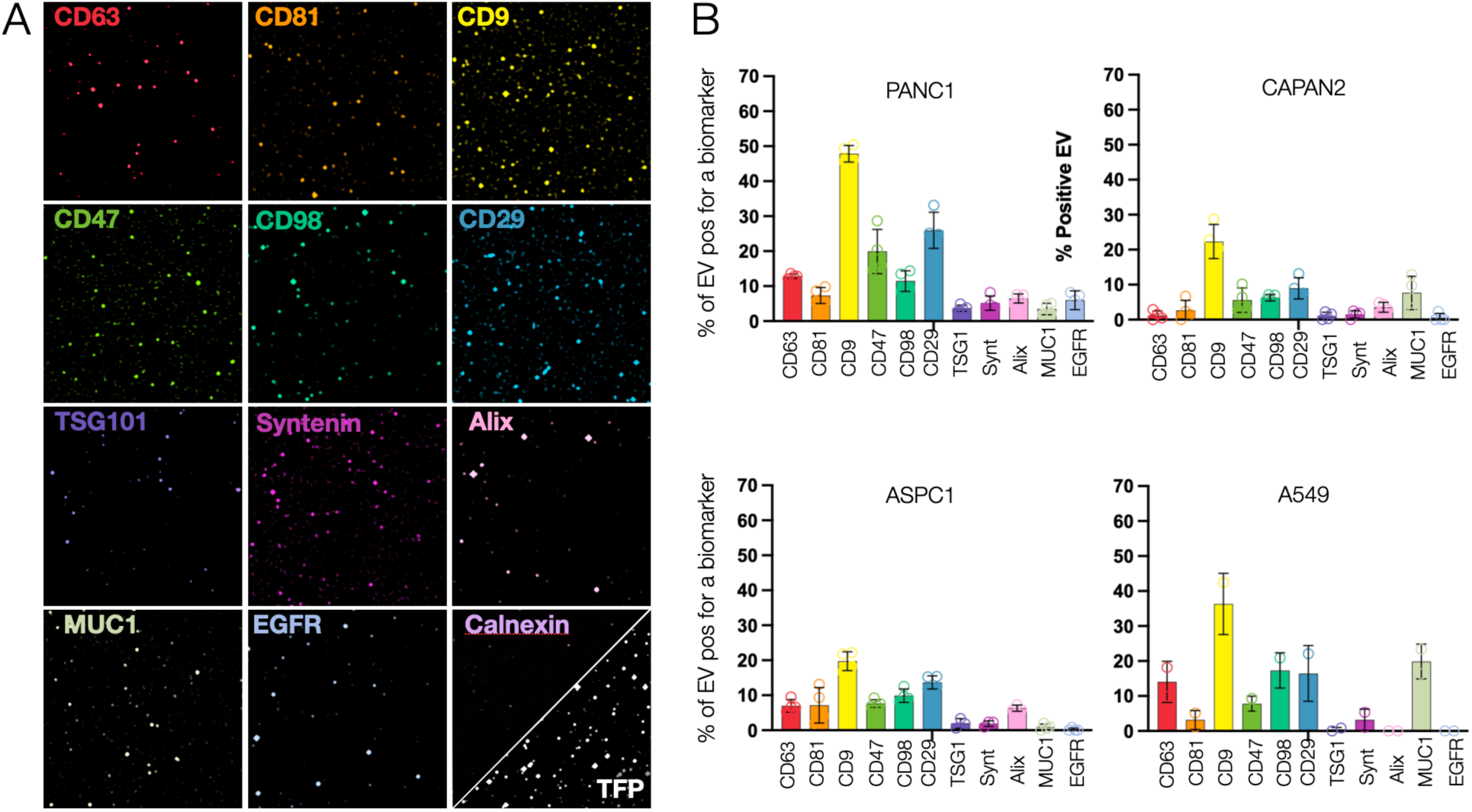
Ubiquitous biomarker analysis in single EV across different cell lines. **A**. High magnification images of EV(obtained from PANC1 cell line) for 12 biomarkers. **B.** Quantitative analysis. Each graph shows the abundance of the indicated biomarker as a percentage of all EV. EV were harvested from PANC-1, CAPAN2, ASPC1, and A549 cells (for other cells, see **Fig. S5-S6**). The most abundant marker was CD9, followed by CD29. There were considerable differences in marker positivity, even for EV from the same origin. For example, in EV from pancreatic cancer cell lines, CD9 ranged from 20 to 50% in vesicles. Calnexin was used as an exclusion marker.

### Concurrence of EV biomarkers on the same EV

We next interrogated the concurrence of different biomarkers in each vesicle. **Fig. 4** shows the distribution of 3 tetraspanins (CD9, CD81, CD63) in PANC-1 and CAPAN-2. Note that 24.8% of all PANC1 EVs and 52.6% of all CAPAN-2 EVs had no tetraspanin. Across four cell lines, PANC1, CAPAN-2, ASPC1, and A549, and average of 39% of EV had no tetraspanins (20-52.6%), 39% had only one tetraspanin (35.6-50.0%), 22% had two (9.4-35.6%), and only 5.4% had all three tetraspanins (2-8.4%).

**Fig. 4:**
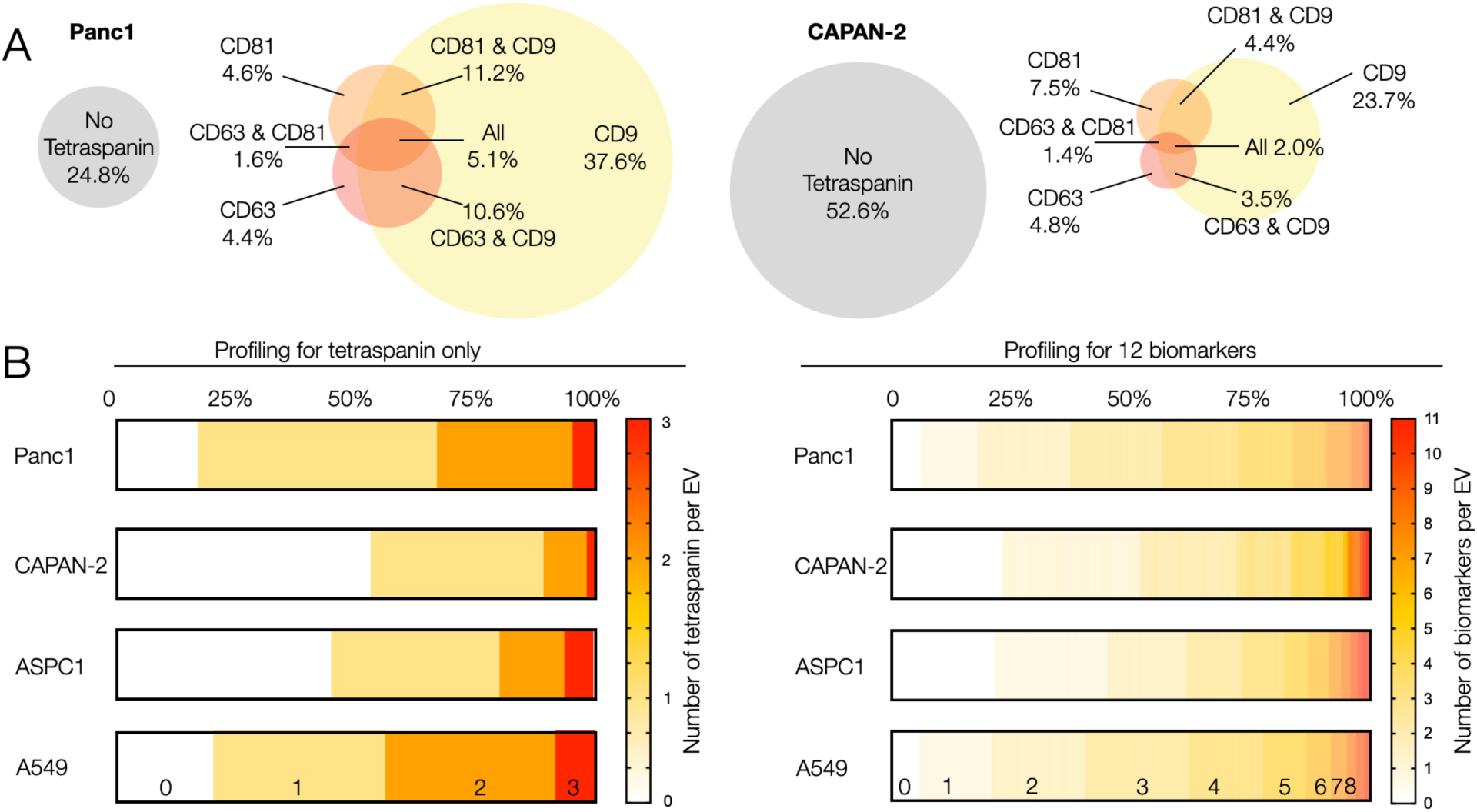
Analysis of biomarker concurrence. **A**. Analysis of tetraspanins (CD9 (TSPAN29), CD63 (TSPAN30) and CD81 (TSPAN28)). Analysis of PANC-1 EV (left) and CAPAN-2 EV (right). Note that 24.8% of all PANC-1 EVs do not express any of the three tetraspanins. In CAPAN-2, 52.6% of EVs do not express any of the three tetraspanins. CD9 was the most common tetraspanin in both EV types. **B**. Analysis of tetraspanin (left) or 12 other biomarkers concurrence (right; see Table S1 for markers) across individual EV obtained from 4 cell lines. Only 2-10% of all EV show all three tetraspanins. Roughly 50% of all vesicles show < 3 of the 12 other biomarkers combined in a given vesicle. Only a small percentage of vesicles shows > 5 of the biomarkers.

We expanded this analysis to all 12 biomarkers tested and across all EV types (**Fig. 4B**). Using TFP_350_ as a ubiquitous EV reference stain, we determined that the most common patterns across PANC1 EV with this panel were 2 or 3 (38%) detectable markers, 4 or 5 biomarkers (27%), 0 or 1 EV biomarkers (18%), followed by 6 or 7 biomarkers (12%). Similar distributions were observed in the other three cell lines. Across the set, only a small percentage of vesicles showed 8 or more simultaneous biomarkers: PANC1: 4.22%; CAPAN-2: 0.26%; ASPC1: 2.75%; A549: 2.00%. Recently identified “ubiquitous” EV biomarkers^33^ were present only in a small fraction of individual single PANC1 EV (e.g., syntenin: 5.1%).

### Oncogene and tumor suppressor protein identification in vesicles

Biomarker constellations and cancer-specific biomarkers (e.g., mutated proteins such as KRAS^mut^ and P53^mut^) have been well established for human cancers. It has also been shown that some of these biomarkers are present in EV, albeit at very low rates in early cancers^19^. Therefore, to use EV diagnostics for early cancer detection, one would like to enrich cancer biomarker-positive EV. There are at least three challenges at hand; i) to separate EV from other circulating vesicles, ii) to separate tumor EV from host cell EV, and iii) to identify EV surface proteins that allow enrichment of cancer protein protein-positive EV (KRAS^mut^ and P53^mut^ are intravesicular proteins). To facilitate such analyses, we performed KRAS^G12D^, KRAS^G12S,^ and KRAS^G12V^ profiling of single EV derived from RAS-positive A549, LS180, and PANC-1 cell lines and asked how common tetraspanins are in oncogene-positive vesicles.

**Fig. 5** summarizes the KRASmut data. We show that KRAS^G12D^ is detectable in 45% of PANC1 EV, in 40% of ASPC1 EV, and in 20% of LS180 EV. KRAS^G12S^ was detectable in 30% of A549EV, and KRAS^G12V^ was detectable in 15% of CAPAN-2 EV. Conversely, mutated KRAS EVs could not be detected in KRAS wild-type EV such as those from A431 and MCF-7 (**Table S2**). We next asked how many KRAS^mut^ positive EV would be missed if one were to perform affinity purification with one or all of the tetraspanin markers. Our results show remarkable EV loss rates, with up to 80% of all oncogene-positive EV being missed with routinely used CD63 affinity purification. This inefficiency is problematic for clinical samples and early cancer detection, where the KRAS^mut^ positivity is < 0.1% of all EV^19^. While a pan-tetraspanin capture strategy can improve detection yield, our results indicate that 35-45% of KRAS^mut^ positive EV would still be missed.

**Fig. 5:**
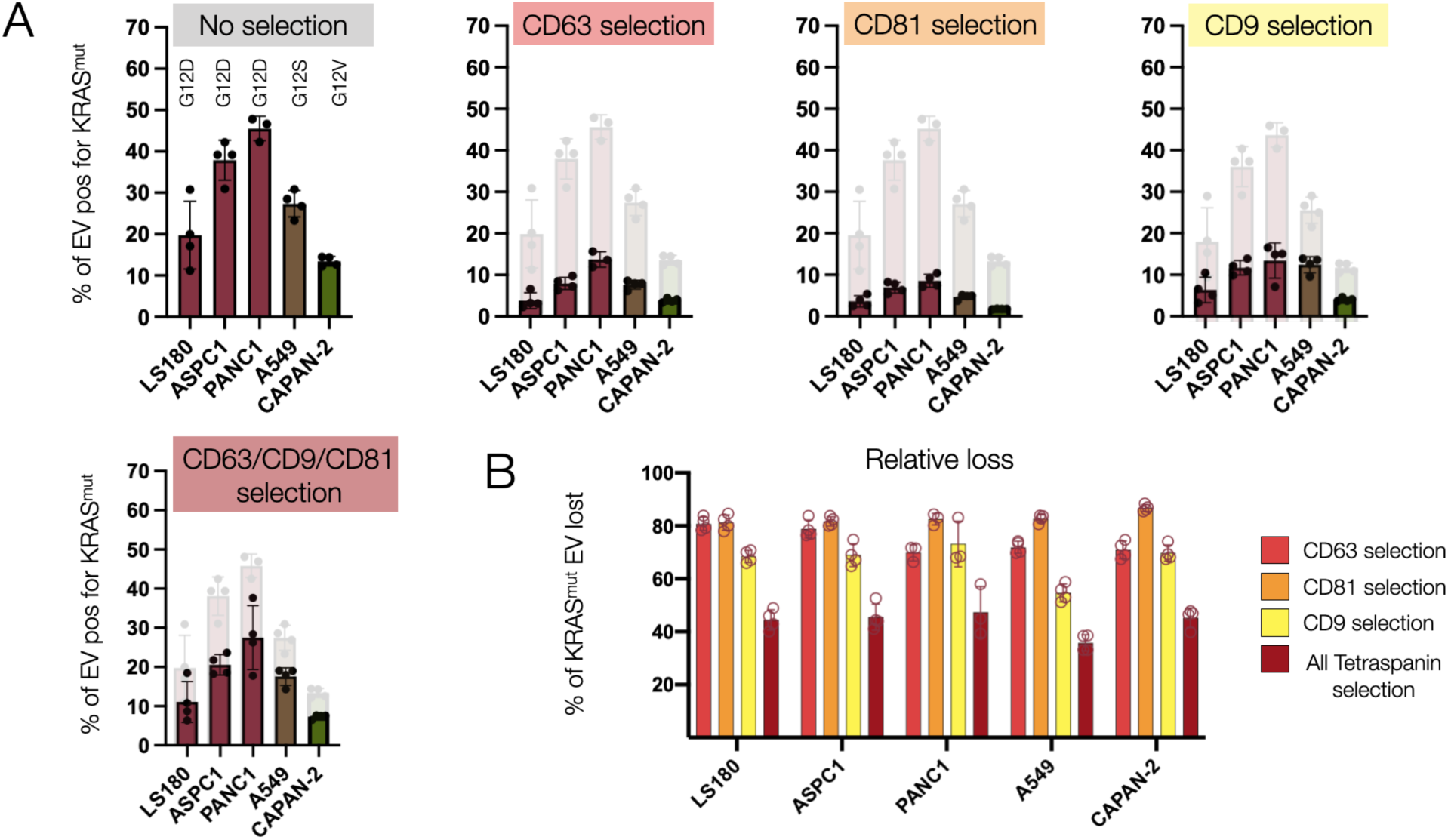
Analysis of EV carrying cancer-specific KRAS mutated proteins. EV carrying cancer-specific mutated proteins (KRAS^G12D^ (PANC1, ASPC1, LS180); KRAS^G12V^ (CAPAN-2); and KRAS^G12S^ (A549) were analyzed for co-expression of tetraspanins commonly used for affinity purification of EV. **A**. Profiling of EV for different KRAS^mut^ proteins in EV. Note that 15-45% of all EV exhibit KRAS^mu^t protein in their EV. This number decreases considerably when accounting for concurrent tetraspanin positivity. **B.** The data show considerable loss (50-80%) of KRAS^mut^ positive EV with affinity purification.

### Size analysis

We next explored whether there was a size dependency on biomarker positivity across vesicles 30-150 nm in size. We first calibrated MASEV size determination against TEM, DLS, and NTA as gold standards. There was a good correlation between the different methods of size measurement summarized in **Fig. S4**. The mean EV size was ∼100 nm (30-150 nm; **Fig. S3**) depending on cell type (**Fig. S4)**. **Fig. S7** summarizes the biomarker analysis in PANC1 as a function of EV size. This experiment shows biomarker expression levels for different EV types (LS180, ASPC1, PANC1, A549, CAPAN-2) in vesicles < 50 nm and > 50 nm as determined by MASEV imaging. On average, there was no significant difference in biomarker expression between smaller and larger EV. The notable exception was for KRAS^mut,^ which appeared at higher amounts in larger EV (> 50 nm). These results were confirmed by electron microscopy using KRAS^G12D^ gold nanoparticles (AuNP). Mutated KRAS was more commonly found in larger EV. Combined, this data suggests that while there is no need for further size fractionation to measure common biomarkers in EV populations 30-150 nm, future study may be warranted to define the optimal size distribution for detection and analysis of key oncogenic markers.

### Mapping EV populations

Having established cyclic molecular analyses of EV, we next set out to map the EV results across different EV types. **Fig. 6A** and **Fig. S8** show representative maps of 12,000 single EV analyzed for 12 biomarkers. To the best of our knowledge, this represents the first large-scale single EV map obtained with expanded multiplexed profiling capabilities. To determine whether expanded multiplexing could resolve different types of EV, we pooled the data for the single EV from four cell lines as a pilot test for determining of tissue of origin and applied dimensionality reduction using t-distributed stochastic neighbor embedding (t-SNE) analysis. We used 3000 single EV from each of the 4 cell lines shown in Fig. S8 and ran tSNE analysis for either the complete biomarker panel or restricted three marker subsets. This was done to compare expanded MASEV profiling to the limited single cycle fluorescence profiling of recently published sEVA methods (**Table S3**)^19^. We show that full biomarker multiplexing allows clear separation of EVs from different cell origins (**Fig 6B**), whereas limited biomarker profiling such as the triad of CD9, CD47, and EGFR depicted here, does not allow clear separation. These results suggest that high-multiplexing methods to resolve different EV types based on molecular signatures that cannot be distinguished with routine spectrally resolved 3/4-color imaging.

**Fig. 6:**
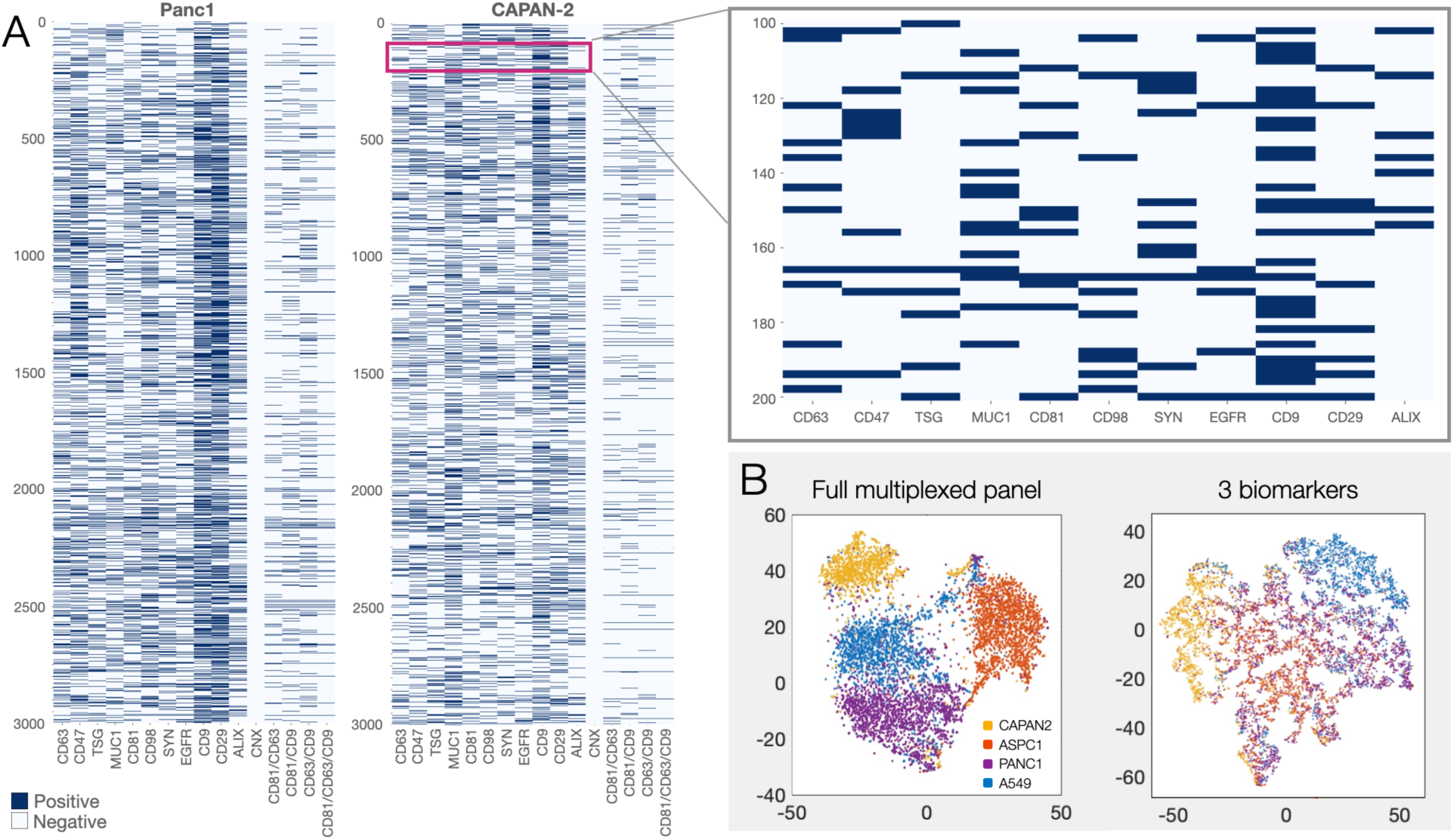
Mapping of thousands of single EV. **A.** Each row represents a single EV and each column a single or combined tetraspanin biomarker. Panc1 EV is shown on the left and CAPAN2 EV on the right (for extended profiling of other cell-line derived EV, see **Fig. S8**). Top right: magnified view shows detailed data on individual EV. No clustering was applied in these examples to illustrate the heterogeneity of biomarkers across the large number of single EV. **B.** Dimensionality reduction of the complete dataset involving 12,000 single EV from 4 different cell lines (yellow: CAPAN2, orange: ASPC1, purple: PANC1 and blue: A549; see Fig S8). Note that full biomarker multiplexing allows clear separation of EVs from different origins. Limited biomarker multiplexing with CD9, CD47, and EGFR, such as done by spectrally resolved 3-color imaging, does not allow clear separation.

## Discussion

EV-related research has grown considerably, with many new technological developments and alternative approaches in isolation and sensing. Yet, there is a surprising scarcity of data related to vesicles’ compositional heterogeneity, and many fundamental questions remain. A better understanding of protein patterns in a single EV will be essential in developing future isolation/capture strategies and determining the proper limits for early cancer detection. Furthermore, a deeper single vesicle analysis will ultimately help understand vesicle heterogeneity and potentially organ-of-origin analysis for vesicles in plasma.

In the current study, we demonstrate the feasibility of performing a highly multiplexed analysis of proteins in single EVs. This analysis was enabled by the recent development of a bioorthogonal, immolating click chemistry that allows rapid cleavage of fluorochromes from antibodies^26, 27^. The fast kinetics and exceptionally complete cleavage allow cycling and repeat staining of individual vesicles in simple microscope slide flow chambers. Using this technology, we profiled and ranked putative biomarker positivity in different model cell lines. We show remarkable heterogeneity of tetraspanins and other EV biomarkers within relatively homogenous cell culture samples. This finding explains the high abundance of such proteins in bulk analyses but scarcity in individual vesicles. This finding has ramifications for future single EV analysis, affinity purification, and detection of rare EV in the early stages of cancer.

Affinity purification remains a standard analytical method to isolate and enrich EV from biological samples. Typical markers include tetraspanins (CD9, CD81, CD63) and other vesicle surface proteins (e.g., EpCAM or EGFR). While such methods allow enrichment of homogenous EV populations, it has been less clear how many EVs with diagnostic potential are lost in such a process. Our profiling data suggests that single tetraspanin purifications can lose up to 80% of KRAS^mut^ EV, and tetraspanin cocktails (e.g., CD9+CD63+CD81) miss 36-47% of KRAS^mut^ positive EV (**Fig. 5**). Since EV with mutated oncoproteins occur in < 0.1% of patient samples with early cancers^19^, such affinity purification may be counterproductive for clinical analysis.

The MASEV method allows broad and deep profiling of individual EV, which has not been possible to date. Since the method is imaging-based, it allows concomitant size analyses and molecular biomarker expression in individual vesicles. Combined, this allows for a deeper characterization of the heterogenous EV population in living systems. We mapped the single EV heterogeneity across vesicles obtained from different cell lines in one potential application, validating methods for future analyses of clinically-derived samples. It is now well understood that most host cells will produce EV and shed them into circulation, from which they are cleared with well-established kinetics^41^. The question is whether deep multiplexing methods such as MASEV developed here will allow an organ-of-origin analysis of circulating vesicles. While this field is currently only nascent, deeper multiplexing will be necessary to enable such analyses, analogous to high-dimensional flow cytometry, in classifying circulating immune cells.

Several single EV analytical methods have previously been described (**Table S3**), but virtually all of them are limited to a few “color channels” during a single round of analysis. In a recent study^19^, we used 3-4 channels (not cycles) for analyzing single EV. We had to perform analysis in aliquoted samples for deeper examination since the multi-cycle method shown here had not yet been developed. The current research indicates that the MASEV method is a vast improvement in multiplexing and performing analyses on simple flow chambers that enable reagent preservation. Although other non-imaging-based single EV technologies are available, most are technically complex (EVseq^21^, ddPCR^20^) or costly. In contradistinction, MASEV is fast and inexpensive, compatible with plasma sample analysis, designed for cancer biomarker analysis, and can be extended to other vesicle types (e.g., microvesicles, tumor-educated platelet vesicles). The next logical step is to use the validated method to analyze clinical samples in prospective well-controlled trials.

## Materials and Methods

### Cells

A431, A549, AsPC-1, CAPAN-2, MIA PaCa-2, PANC-1, and LS180 were obtained from ATCC (Manassas, VA). Cells were grown in Dulbecco’s modified Eagle medium (DMEM) supplemented with 10% fetal bovine serum (FBS). The medium was changed every 1 to 2 days, and cells were passaged before confluency. After each cell line reached confluency, the medium was collected every other day for seven days for EV isolation. All cells were characterized by IHC and flow cytometry^19^.

### Antibodies

Commercially-available antibodies (**Table S1**) were purchased carrier-free for in-lab modification with Dye-C_2_TCO-NHS linker SAFE probes. Probes were synthesized and activated according to^27^ and stored at -80 °C until use. Antibodies were exchanged into 0.1 M PBS-bicarbonate buffer (pH 8.4) using a 40 k Zeba column (87765 Thermo Fisher) and incubated with a 5- to 12-fold molar excess of the SAFE probe with 10% DMSO for 25 minutes at room temperature in the dark. After conjugation reaction, unbound probes were removed by another two 40 k Zeba columns equilibrated with PBS. To determine the degree of labeling (DOL), the absorbance spectrum of the C_2_TCO-labeled antibody was measured using a Nanodrop 1000 (Thermo Scientific). Labeled antibodies (DOL 2-3) were stored in the dark at 4 °C in PBS until use. Hk-Tz scissors used for cleaving C_2_TCO were prepared according to^26^ and stored at -80 °C until use. Before de-staining, Hk-Tz aliquots were thawed and diluted into PBS up to 50µM.

### EV isolation

We first compared different EV isolation methods to determine which ones would be practical for a clinical workflow (small sample volume, high throughput, fast separation, and low cost). Specifically, EV obtained from i) cell line supernatant or pooled plasma and) were isolated using the following methods: ultracentrifugation, qEV IZON size exclusion chromatography, dual-mode chromatography (DMC)^17^ and enhanced dual-mode chromatography (eDMC)^30^.

Ultracentrifugation was performed as previously described^19, 42^. Briefly, for cell line supernatant, we used ∼160 mL conditioned media from cells cultured at least 48 hrs in complete media containing exosome-depleted FBS (Thermo, A2720803); for pooled plasma, we used 3 mL of plasma diluted to 30-35 mL total in PBS. Two ultracentrifugation steps were performed at 100,000 x g for 70 min to obtain EV pellets resuspended in ∼100 µL PBS. IZON size exclusion using qEV single columns (SP2, IZON Science) was utilized according to manufacturers’ instructions. Briefly, 100 µL of pooled plasma was centrifuged at 1500 x g for 10 min at 4C to remove cellular debris. The supernatant was transferred to a clean tube and centrifuged at 10,000 x g for 10 min at 4C to remove larger particles. The qEV single column was flushed with 4 mL 0.22 µm filtered PBS using an IZON automatic fraction collector. Following column flushing, plasma was loaded onto the column. As soon as the sample entered the column resin, PBS was added, and a 1 mL void volume was collected. Three 0.2 mL fractions containing EV were collected and subsequently concentrated using an Amicon Ultra-4 10 kDa filter (Millipore UFC801024).

DMC^17^ and eDMC^30^ EV purification were done according to the methods of each respective publication. Briefly, size exclusion columns were prepared with an 11 µm nylon net filter (NY1102500, Millipore) placed in the bottom of a 10 mL syringe (BD 302995). 2 mL washed Fractogel EMD SO3-(M) (Millipore Sigma, 1168820100) was first layered in the syringe, followed by 10 mL washed Sepharose CL-4B (Millipore Sigma, GE17-0150-01). Columns settled overnight at 4C for at least 24 hrs before use. Columns were flushed with 10 mL 0.22 µm filtered PBS (pH 6.4 for eDMC and pH 7.4 for DMC) before loading with 500 µL pre-cleared (prepared as above for IZON) pooled plasma. After samples entered the resin bed, PBS was added, and a 4 mL void volume was collected, followed by a 2 mL EV-containing fraction. EV were concentrated using an Amicon 10 kDa filter, and the eDMC sample was further buffer exchanged by the addition of PBS pH 7.4 and repeat centrifugation.

## Validation

### Antibody validation

Antibodies are all commercially available and were selected based on the availability of rigorous validation data by Western blotting or flow cytometry (Table S1). All antibodies were further validated in-house by Western blot using cells with known target expression. Antibodies were validated by Western blot or immunofluorescence on EV using nonspecific IgG as a negative control^43–45^.

### Permeabilization protocols

Several EV biomarkers are intravesicular and thus necessitate semi-permeabilization to expose targets to labeling antibodies. We experimented with different permeabilizers (Triton X-100, Tween, SDS), concentrations, and timing protocols. We adopted the use of a 0.001% Triton X-100 (X100, Sigma-Aldrich) solution for 15 min at RT as it resulted in the highest labeling efficiency without destroying channel-bound EV.

### Quality of EV purification

The quality of purified EV methods was compared by Qubit protein analysis (Thermo, Q33211), NanoSight nanoparticle tracking analysis (Malvern), and Western blot for TSG101 (GeneTex GTX 70255), CD63 (Ancell 215-820), ApoB100 (R&D MAB4124) and ApoA1 (R&D MAB36641), sEVA^19^ and electron microscopy.

### Chemical purity

The purity of antibody dye-C_2_TCO-NHS linker SAFE probes, HK-Tz scissors, and TFP_350_ were determined by mass spectrometry and chemical NMR.

### Pan-EV staining with fluorescent TFP

To determine biomarker positivity as a fraction of total EV, we used the TFP protein labeling methods to stain for all EV. This method has been well characterized^19^ and is superior to DiO, DiI, and other lipid staining methods. In brief, the AF350-PEG_12_-TFP conjugate was prepared as described previously^19^. In most cases, 500 ng of EV was combined with 1 µL of Dye-PEG_12_-TFP. This ratio was scalable up to 1500 ng of EV input. The EV and AF350-PEG_12_-TFP reactions were brought to a final volume of 14 µL with filtered PBS and incubated in 1.5-ml Eppendorf tubes protected from light and under agitation using HulaMixer for 2 hours at room temperature. Excess AF350-PEG_12_-TFP was removed using Zeba Micro Spin Desalting Columns, 40K MWCO, 75 µL according to the manufacturer’s protocol.

### Device fabrication and assembly

Glass slides and cover glass (cut to size with a diamond scribe) were prepared as described with slight modifications^29^ (one round of KOH treatment and adding a final isopropanol rinse). First, the glass was extensively cleaned by sonicating in Sparkleen detergent (Fischer Scientific) and acetone baths, followed by rinsing with Milli-Q water. Next, Glass slides were rinsed with isopropanol, dried with N_2_ gas, and stored under a vacuum to prevent dust accumulation. Cover glasses were then further sonicated in a 2M KOH bath for 1 h to activate the glass, rinsed with isopropanol, and thoroughly dried with N_2_ gas to remove all water. To form a hydrophobic silane layer on the coverglass, activated glass was incubated with 50 µL fresh dichloro dimethyl silane (DDS) (440272, Sigma) solution in 75ml hexane for 1.5 h, rinsed and sonicated with hexane, rinsed with isopropanol, and thoroughly dried under N_2_. The treated cover glasses were stored in a vacuum desiccating chamber until used. The coverglass surface was maximally activated for subsequent attachment of EV.

Microfluidic devices were prepared following a recent description of an inexpensive device for cell staining^28^. Channels were cut from 50 µm thick double-sided 3M VHB adhesive tape (F9460PC) using a Silhouette Cameo 4 craft cutting tool (Silhouette USA). Cut tape layers were adhered to a glass slide and enclosed with a treated cover glass. The hydrophobic coverglass prevented the device from ‘wetting out’ over multiple staining and destaining cycles. After sealing the chamber, EVs were flown into the channel and incubated at RT for 30 minutes, then incubated with 0.05% Tween-20 solution (003005, Thermo-Fischer) for 15 min at RT to passivate the remaining coverglass surface. Tween-20 solution was prepared in buffer as 10 mM Tris, 50 mM NaCl, pH 8.0. EVs were next permeabilized by flowing through 0.001% Triton X-100 (X100, Sigma-Aldrich) in PBS and incubating for 15 minutes. Finally, SuperBlock (37580, Thermo Scientific) was flown through the device and incubated for 30 minutes to passivate the untreated glass slide base.

### Cyclic EV profiling (MASEV)

EVs were profiled by cyclically labeling and cleaving off fluorochromes from Ab-C_2_TCO-Fl probes. To label TFP-stained EVs, probes were diluted to 10 µg/ml in SuperBlock and passively pumped through the devices. The device was then incubated at RT for 1 h in a humid enclosure to prevent evaporation. Excess probes were rinsed with PBS (5 µL three times). After imaging, labels were cleaved off by flowing through 50 µM of HK-Tz scissors (5 µL three times) and incubating for 10 minutes at RT. Scissors were rinsed with PBS (5 µL three times) to prepare the device for subsequent cycles. Each labeling round contained three spectrally distinct probes (MB488, AF555, AF647). Pumping was done by pipetting liquid into one of the two evacuated reservoirs of the device, whereby the small-radius droplet generated differential Laplacian pressure to drive fluid flow (video S1).

### Image acquisition

While obtaining EV images with super-resolution or confocal microscopy is possible, we opted to use a more commonly available multichannel epifluorescence microscope typically used for multiplexed cell imaging^46^. Specifically, images were acquired on an upright Olympus BX63 microscope using 20-40x objectives. EVs were brought into sharp focus with the DAPI filter set to image the universal and non-cleavable 350-TFP stain. Images were acquired with a 4000 ms long exposure. Proper focusing is critical since lower-signal nano-sized EVs can be lost even with minor changes in z-plane settings. The live image display was monitored to ensure ideal focusing by looking at the smaller/dimmer EVs with 2x2 binning and manually adjusting until a minimum pixel radius was observed. Suitable FOVs were kept near the center of each channel to avoid uneven background from near-perimeter areas (e.g., scattering, autofluorescence from tape). After obtaining the total EV-TFP image, the samples were imaged for 4000ms with the FITC, Texas Red, and CY5 filter sets.

### Image processing and analysis

Image analysis was completed using ImageJ (NIH) and Python v3.7.0 (**Fig. S9**). Images were aligned, background-subtracted, and segmented, and the average-fluorescence intensity was measured. Images were aligned using phase cross-correlation to correct for translations that occurred from removing and replacing the flow cell under the microscope between cycles. A region-of-interest (ROI) mask was created by thresholding the average intensity of TFP-labeled EVs with the Triangle algorithm^47^.

This way, the number of TFP-identifiable EV per FOV, average EV size, and therefore total percent coverage of EV could be standardized across images in line with the standardized amount of EV deposited per slide. All EV were identified by TFP350 labeling, and EV with an area of fewer than 5 pixels were excluded from the analysis. This mask was applied to the background-subtracted 488 nm, 555 nm, and 647 nm channels to measure EV signals iteratively.

### Electron microscopy

To compare MASEV staining to an accepted gold standard, we used comparative immunogold labeling for electron microscopy (EM)^15, 48^. These studies were performed in EV harvested from KRAS^G12D^ positive AsPC1 cells. Aliquoted samples were processed for EM (MUC1-5nmAuNP, KRAS^G12D^-10nmAuNP, EGFR-15nmAuNP) and MASEV (CD63-MB488, CD81-AF555, CD9-AF647, KRAS^G12V^-MB488, KRAS^G12D^-AF594, KRAS^G12S^-AF647). Data were used for i) size determination and ii) biomarker comparison.

EV were pelleted for fixation and ultrathin cryosectioning by ultracentrifugation of IZON size-exclusion purified EV (as described above). PBS was removed from the ultracentrifuge tube, and 4% paraformaldehyde in PBS (pH 7.4) was gently overlaid on the EV pellet for 2 hrs at room temperature. After 2 hrs, PFA was carefully removed and replaced with PBS. The EV pellet was then incubated in 2.3 M sucrose and 0.2 M glycine in PBS for 15 min, followed by freezing in liquid nitrogen. The frozen sample was sectioned at -120°C, and 60-80 nm sections were transferred to formvar-carbon coated copper grids. Immunogold labeling was done at room temperature on a piece of parafilm. Grids were floated on drops of 1% BSA for 10 minutes to block nonspecific labeling, were then transferred to 5 µl drops of primary antibody and incubated for 30 minutes, washed in 4 drops of PBS (total 10 min) before incubation in Protein A-gold (5nm) for 20 min. Grids were washed in 2 drops of PBS followed by 4 drops of water (total of 15 min). The labeled sections were contrasted and embedded in methylcellulose by floating the grids on a mixture of 0.3% uranyl acetate in 2% methylcellulose for 5 minutes. Excess liquid was blotted off on a filter paper, and the grids were examined in a JEOL 1200EX Transmission electron microscope (JEOL, 11 Dearborn Rd, Peabody, MA 01960). Images were recorded with an AMT 2k CCD camera. (Advanced Microscopy Techniques Corp., 242 West Cummings Park, Woburn, MA 01801 USA).

## Acknowledgments

We thank Dr. Hakho Lee for many helpful discussions and Drs. Hannes Mikula and Martin Wilkovitsch for synthesis of fluorescent C2TCO probes. Dr. Kilean Lucas assisted with designing the fluidic chamber and performing early feasibility experiments. This work was partly supported by the following NIH grants: R21CA236561 and P01CA069246. JS, SF, and HP were supported by T32CA079443.

## Contributions

Conceptualization: RW

Methodology: all authors

Data curation: JS, SF

Validation: all authors

Computation: HP, SF, JS

Supervision: RW, JCTC

Writing - original draft: RW

Writing - review, and editing: all coauthors

Funding acquisition: RW

Resources and administration: RW

## Conflict of interest

RW is a consultant to ModeRNA, Lumicell, Seer Biosciences, Earli, and Accure Health. HP is a consultant to Aikili Biosystems. JS, SF, and JCTC report no industrial interactions.

## Supplemental information

**Fig. S1:**
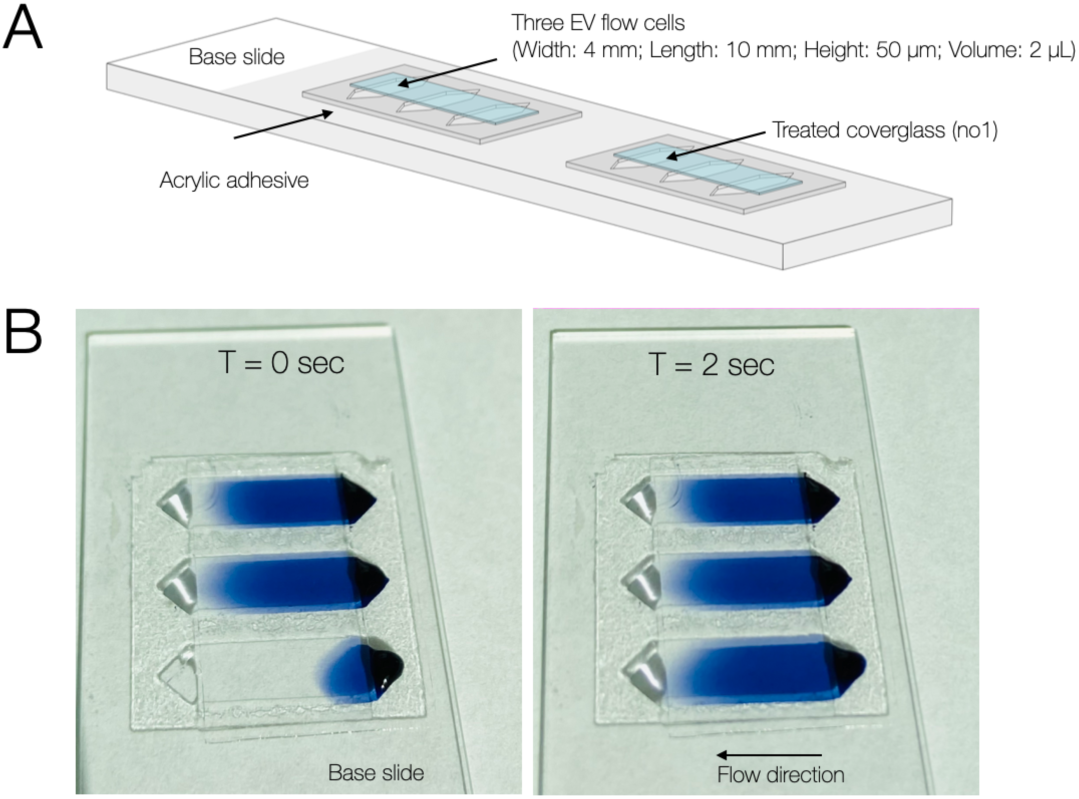
Slide set-up. A. Multiple flow-cell construction on a microscope slide. The flow cells are assembled from acrylic adhesive cut into multiple channels and sandwiched between microscope slides and cover glass. The footprint of the adhesive ( ∼22 mm x 24 mm ) accommodates multiple channels per single base slide. The shape of the pressure-sensitive adhesive cut-out (width 4 mm length 12 mm height 50 µm) allows pump-free flushing driven via Laplacian pressure^28^. Using this design, flow rates of ∼1 µL/s could be achieved within the 2 µL channel. The hydrophobic silanization treatment adheres to EV and prevents the flowcell reservoir from leaking or wetting out over the duration of multiple staining rounds. **B:** Staining and washing. For demonstration purposes, a 5µL drop of trypan-blue was placed in reservoirs and shown to rapidly flush the PBS-filled channel displacing PBS into the reservoir (left versus right panels). Using this method, EV adhered to the glass can be efficiently washed, stained, and de-stained within minutes. **See Movie 1** for animation.

**Fig. S2:**
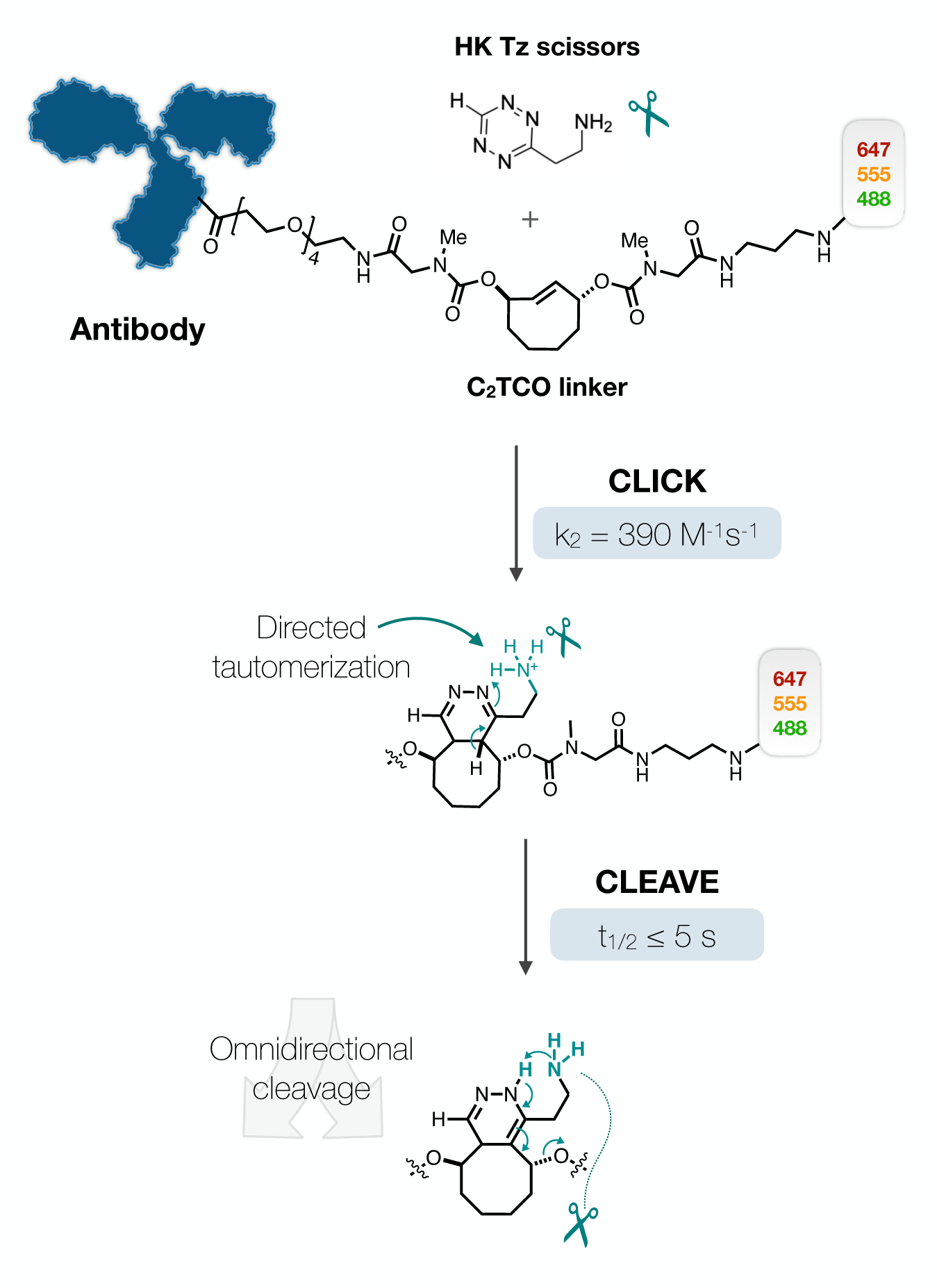
Chemistry of immolative click-to-release linker. Antibodies were labeled with a C_2_TCO linker conjugated to one of three fluorescent dyes: MB488, AF555, or AF647. The fast click reaction between C_2_TCO and the HK-Tz s scissors has a rate constant of 390 M^-1^s^-1^, enabling complete reaction within 4 minutes at 50 µM Tz concentration. The HK ammonium side-chain (NH3^+^ at physiologic pH) directs tautomerization to the direct formation of a rapidly-releasing isomer irrespective of click orientation (oriented left or right in the diagram). Participation of the side chain in the subsequent cascade elimination achieves quantitative release with a cleavage half-life

**Fig. S3:**
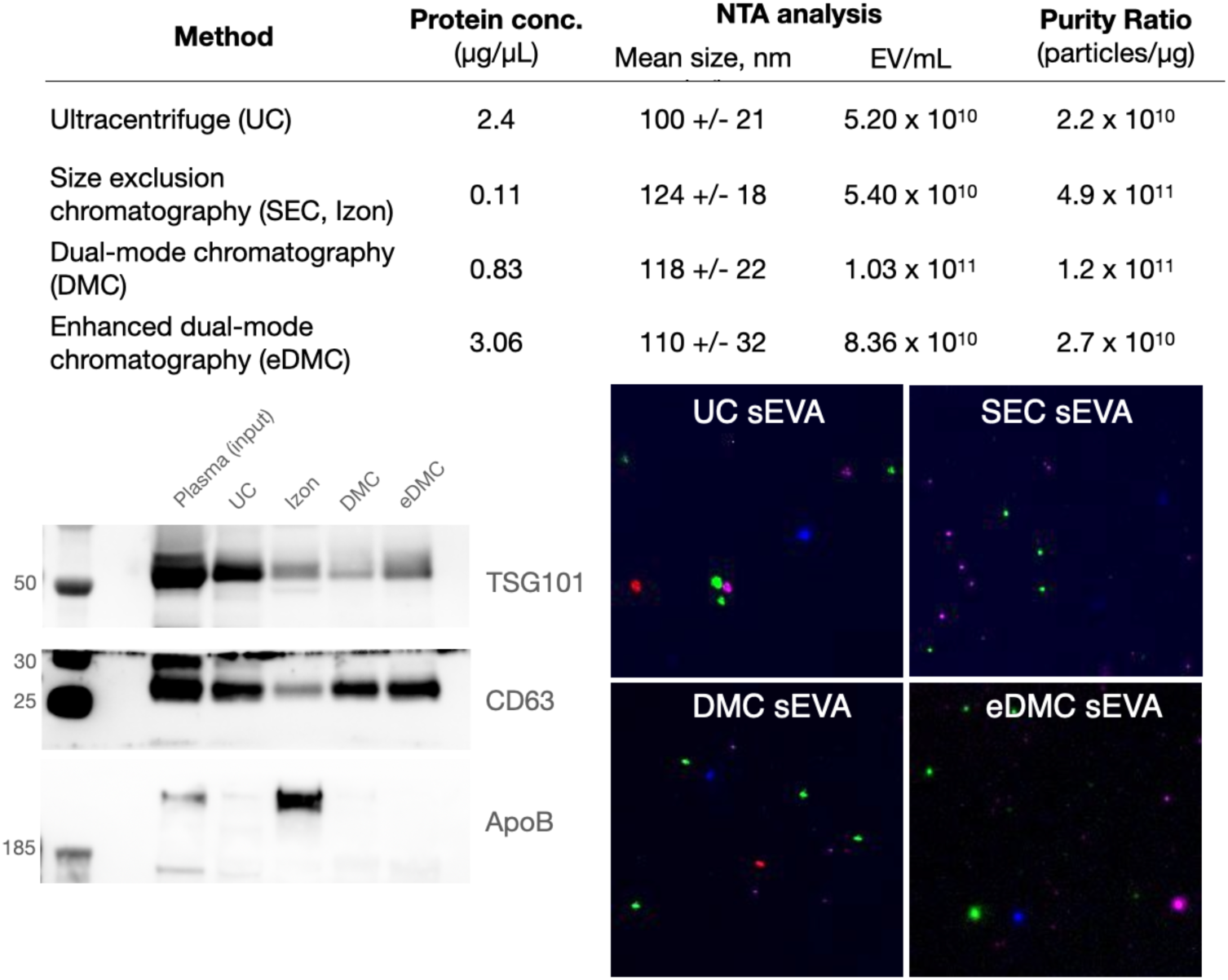
Pre-analytical purification. EV were purified by ultracentrifugation, size exclusion chromatography (IZON qEV single), dual-mode chromatography^17^, and enhanced dual-mode chromatography^30^. EV were compared for total protein concentration (Qubit), nanoparticle tracking analysis (Nanosight), Western blot (ApoB100, TSG101, CD63), and MASEV analysis. Note that advanced size exclusion chromatography methods (DMC, eDMAC) generate pure EV populations combining simplicity, speed, and translatability. All images have the same brightness and contrast for comparison mages: TPF: blue; MUC1: green; KRASmut: magenta; EGFR: red.

**Fig. S4:**
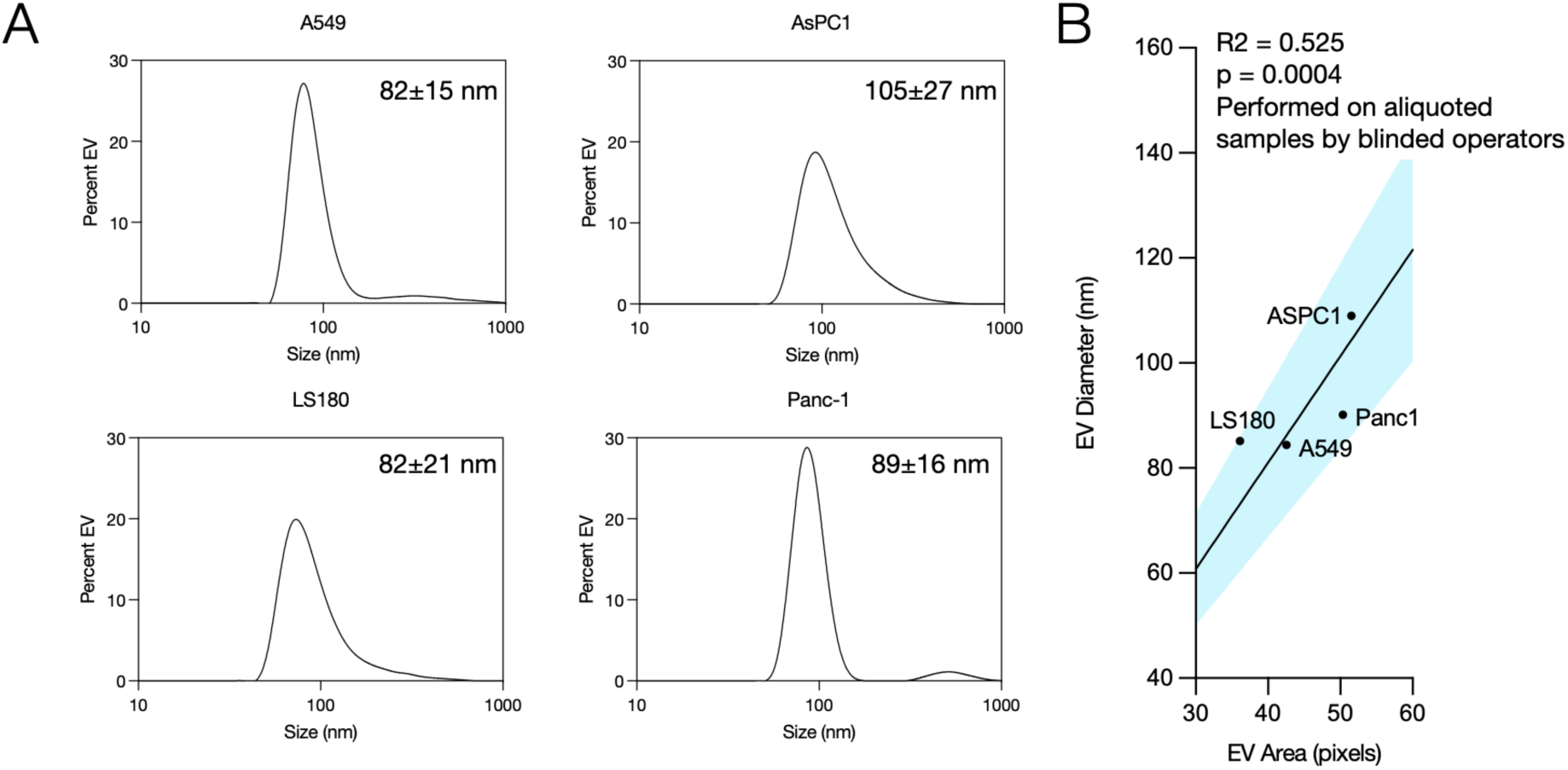
Comparison of size analysis between MASEV and dynamic light scattering (DLS). **A**. DLS graphs of EV samples obtained by eDMC for indicated cell lines. Note the relatively homogenous distribution with mean sizes ∼80-110 nm. **B**. correlation between DLS measurements and MASEV size analysis by imaging (p=0.0004; R^2^ =0.525). The measurements were performed by different operators on different days and who were blinded to samples. There was an even better correlation between MASEV and TEM analysis performed for ASPC1 aliquoted samples (R^2^ = 0.85).

**Fig. S5:**
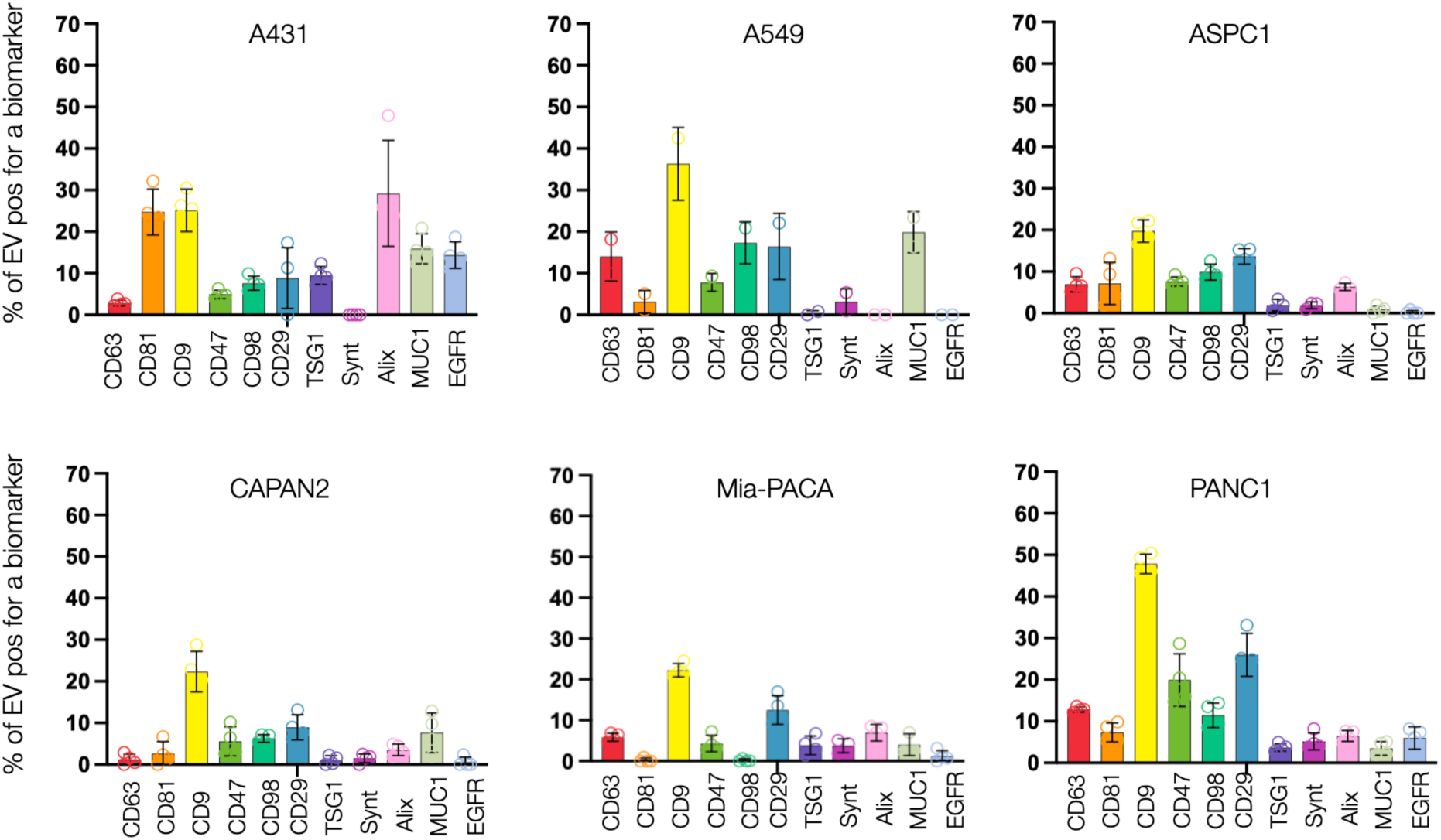
Expansion of MASEV to other cell lines. Summary of EV profiling obtained from 6 cell lines (A431, A459, ASPC1, CAPAN-2, Mia-PACA, and PANC1). The number of EV positive for a given biomarker is indicated on the y-axis.

**Fig. S6:**
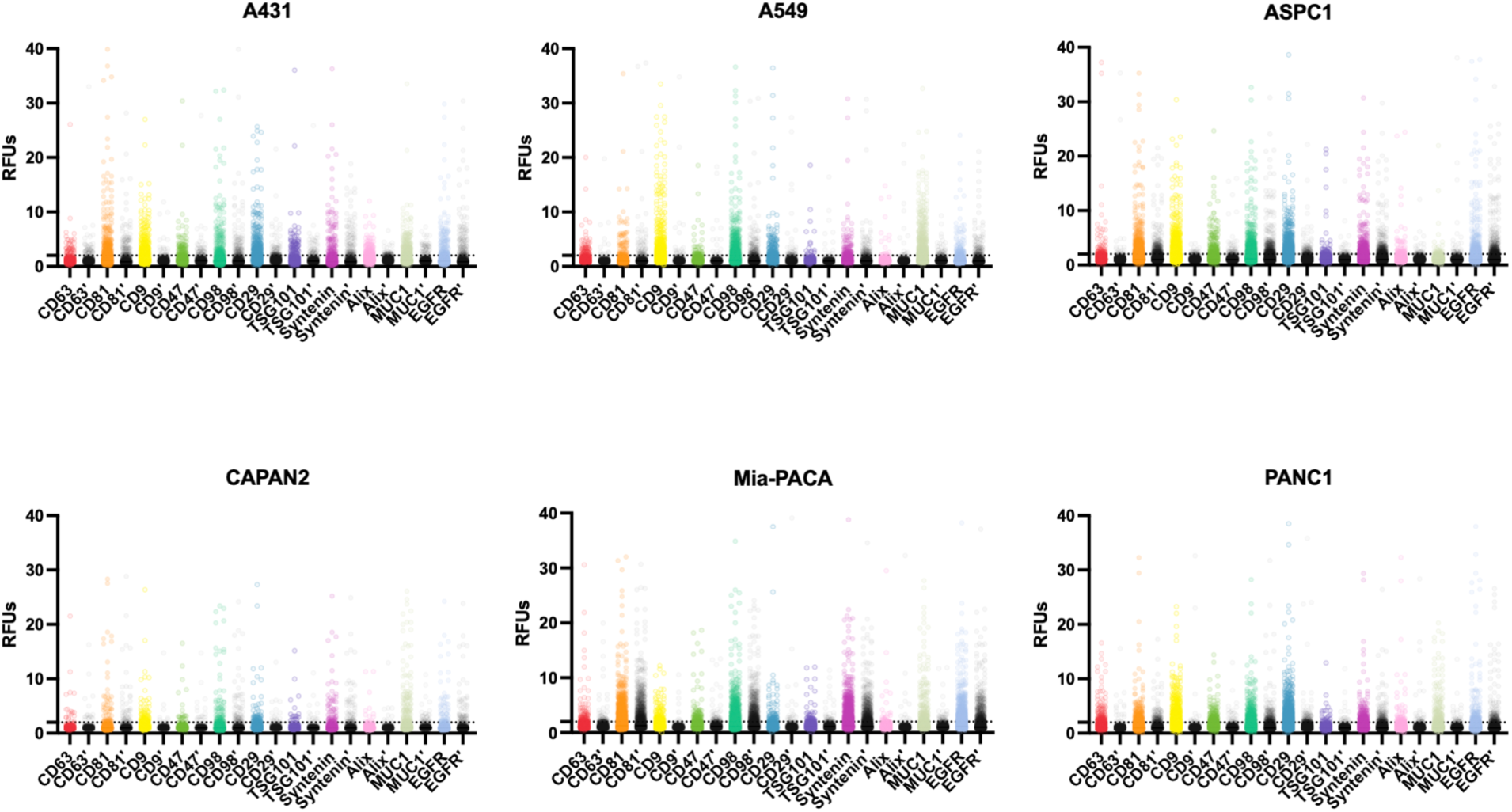
Detailed MASEV analysis. Summary of detailed EV profiling obtained from 6 cell types (A431, A459, ASPC1, CAPAN-2, Mia-PACA, and PANC1). For each of the cell line-derived EV are shown the fluorescence brightness of labeled and subsequently cut (indicated by a prime signal) fluorescence. The dashed line represents background levels.

**Fig. S7:**
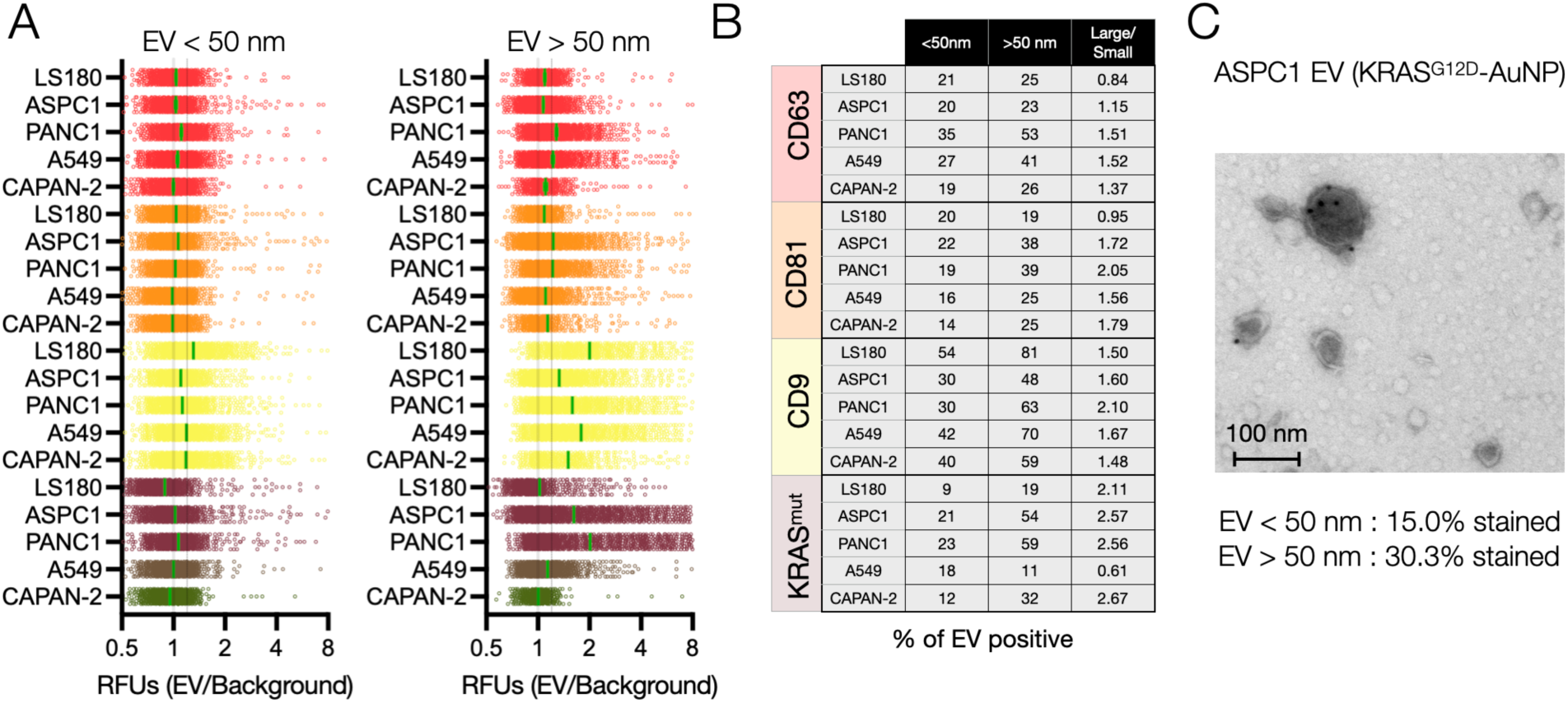
Effect of EV size on biomarker expression. EV were purified by different means, and their size was determined by TEM, DLS, NTA, and MASEV, showing a good correlation (see **Fig. S2-S3**) and a mean size of ∼100 nm (30-150 nm; **Fig. S3**). **A**. This experiment shows biomarker expression levels for different EV types (LS180, ASPC1, PANC1, A549, CAPAN-2) in vesicles < 50 nm and > 50 nm as determined by MASEV imaging. B. Tabulation of results. On average, there was no significant difference between smaller and larger EV, except KRASmut, which appeared at slightly higher amounts in larger EV. **C**. Confirmation by TEM using KRAS^G12D^ gold nanoparticles (AuNP).

**Fig. S8:**
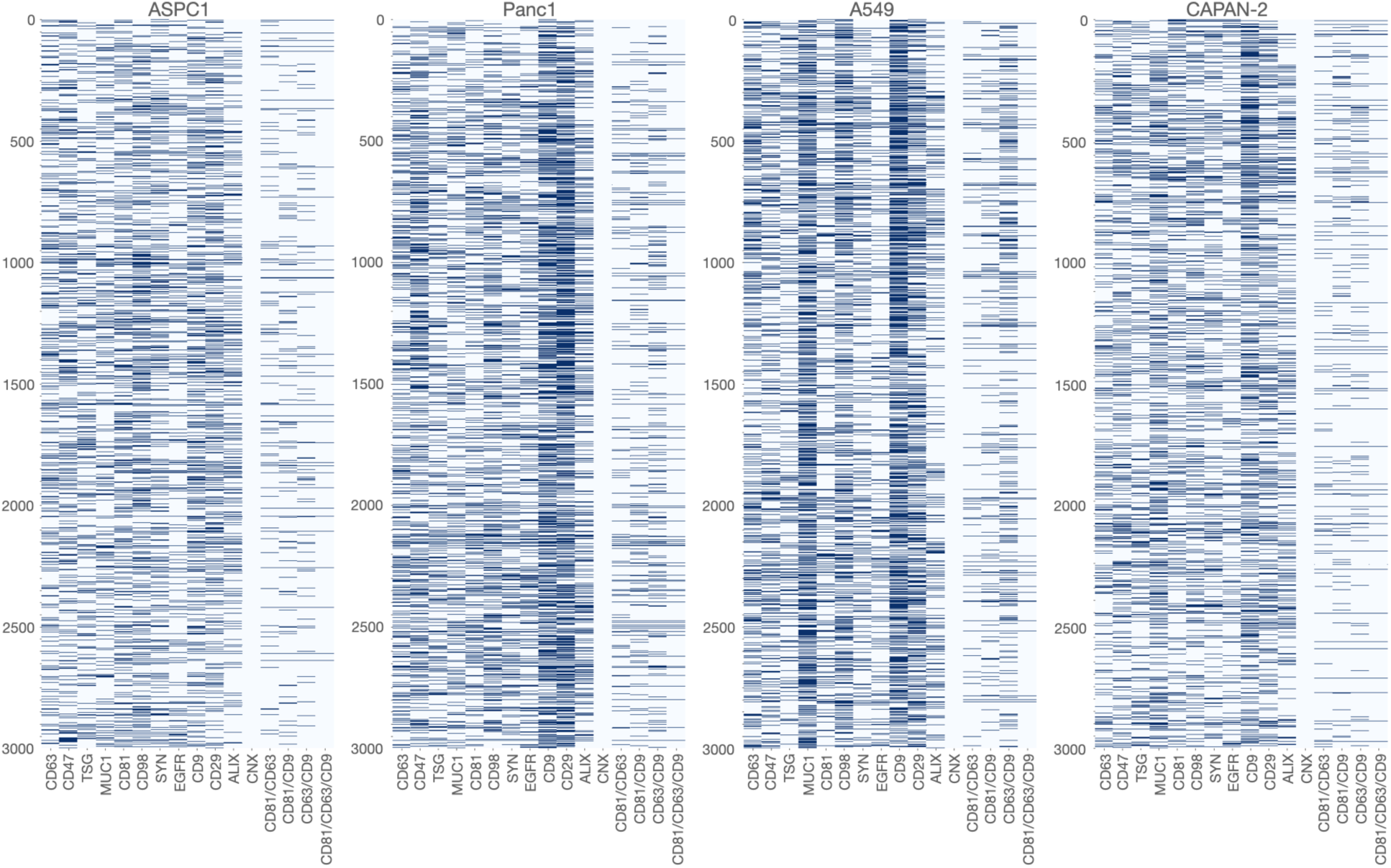
Mapping of 12,000 single EV across 4 cell lines. Shown are biomarker positive or negative single EV. To the left of each plot are single biomarkers, and tetraspanin combinations to the right. No clustering was applied to show the heterogeneity of biomarkers across the large number of EV.

**Fig. S9:**
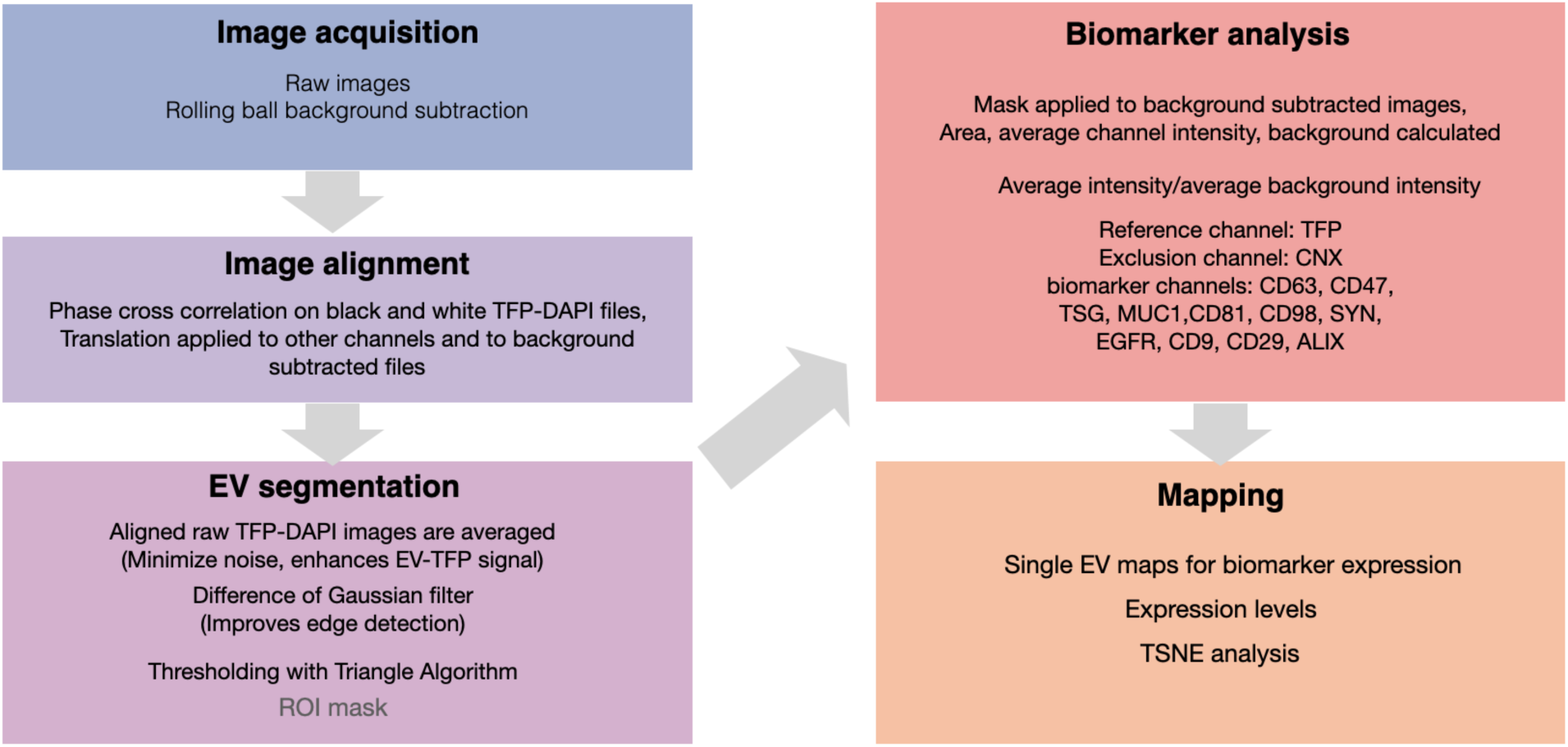
Computational pipeline. Image analysis was performed primarily in Python v3.7.0. Briefly, MASEV images were aligned, background-subtracted, and segmented, and the average-fluorescence intensity was measured. Images were aligned using phase cross-correlation to correct for translations that occur in imaging between cycles. A region-of-interest (ROI) mask was created by thresholding the average intensity of TFP-labeled EVs with the Triangle algorithm. This way, the number of TFP-identifiable EV per FOV, average EV size, and therefore total percent coverage of EV could be standardized across images in line with the standardized amount of EV deposited per slide. All EV were identified by TFP350 labeling, and EV with an area of fewer than 5 pixels were excluded from the analysis. This mask was applied to the background-subtracted 488 nm, 555 nm, and 647 nm channels to measure EV signals iteratively.

**Table S1:**
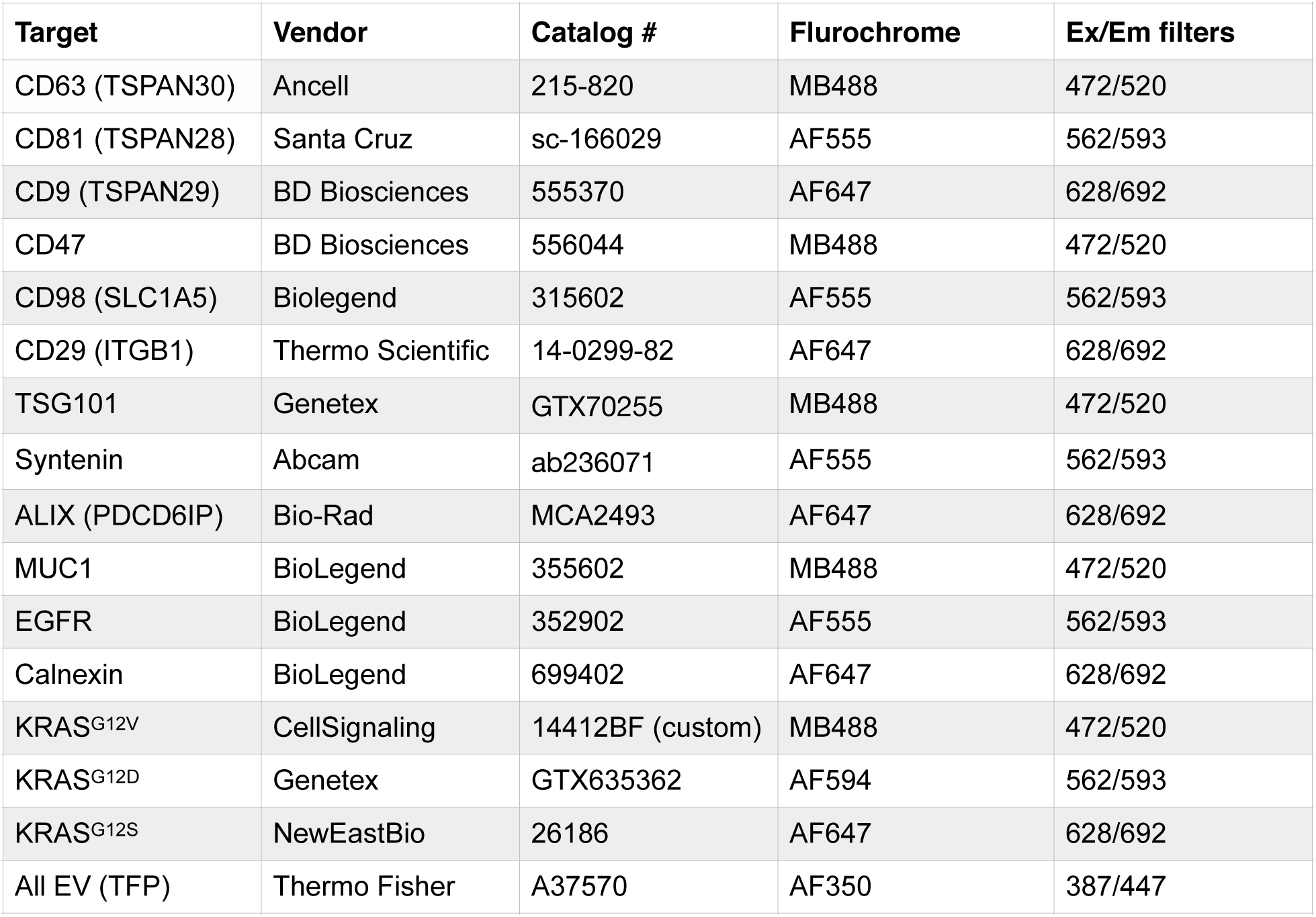
Antibodies. Summary of commercially available antibodies modified with cleavable SAFE linkers and fluorochromes.

**Table S2:** Cell lines used.

| **Cell line** | **Source** | **Sex/Age** | **Site** | **KRAS** | **P53** | **Comment** |
| --- | --- | --- | --- | --- | --- | --- |
| AsPC-1 | ATCC | F/62 | Ascites | G12D | Mut | CRL-1682 |
| PANC-1 | ATCC | M/56 | Primary | G12D | Mut | CRL-1469 |
| CAPAN-2 | ATCC | M/56 | Primary | G12V | WT | HTB80 |
| MIA PaCa-2 | ATCC | M/65 | Primary | G12C | Mut | CRL-1420 |
| A431 | ATCC | F/85 | Skin | WT | None | CRL-1555 |
| A549 | ATCC | M/58 | Lung | G12S | WT | CCL-185 |
| LS180 | ATCC | F/58 | Colorectal | G12D | WT | CL187 |

**Table S3:** Comparison of single EV methods. Summary of different single EV methods to highlight differences in applications and performance.

|  | SEA | ddPCR | EV-seq | sEVA | MASEV |
| --- | --- | --- | --- | --- | --- |
| Reference | *ACS Nano* **12**, 494–503 (2018). | *Adv Biosyst* **4**, e1900307 (2020). | *ACS Nano* **15**, 5631–5638 (2021). | *Sci Adv.* 2022; 8(16) eabm3453 | This study |
| Principle | Labeling on glass | Droplet PCR | Droplet sequencing | Labeling in solution, additional purification | Highly multiplexed analysis |
| Low multiplexing (2-5 markers) | Yes | Yes | Yes | Yes | Yes |
| **High multiplexing (> 10 markers)** | No | No | Potentially | No | **Yes** |
| Complexity | + | +++ | +++ | - | - |
| **Loss of EV** | **>95%** |  |  | <10% | **<1%** |
| Staining quality | Dim<br>SNR ~3 | NA | NA | Bright<br>SNR >5 | Bright<br>SNR >5 |
| Use for rare EV | Not suitable |  |  | Suitable | Suitable |
| Clinical (cost, time, robustness) | - | - | - | +++ | +++ |
| Comment | Historic method | TBD | TBD | Practical | Practical |

**Movie 1: EV flow-cell allows rapid staining and rinsing**. **A**. Flow-cell reservoirs are filled with a 5 µL drop of trypan-blue to load the channel or PBS to flush the channel. The hydrophobic silanized coverglass treatment pins the droplet in place and prevents the reservoir from wetting out onto the coverglass. Applying a vacuum to the distal reservoir allows complete droplet loading into the channel. Trypan blue is visibly loaded after one round and rinsed out with PBS after two rounds (5-10 µL). We, therefore, conservatively fill and flush the device three times (15 µL) in all experiments.

